# Polylactide Degradation Activates Immune Cells by Metabolic Reprogramming

**DOI:** 10.1101/2022.09.22.509105

**Authors:** Chima V. Maduka, Mohammed Alhaj, Evran Ural, Michael O. Habeeb, Maxwell M. Kuhnert, Kylie Smith, Ashley V. Makela, Hunter Pope, Shoue Chen, Jeremy M. Hix, Christiane L. Mallett, Seock-Jin Chung, Maxwell Hakun, Anthony Tundo, Kurt R. Zinn, Kurt D. Hankenson, Stuart B. Goodman, Ramani Narayan, Christopher H. Contag

## Abstract

Polylactide (PLA) is the most widely utilized biopolymer in medicine. However, chronic inflammation and excessive fibrosis resulting from its degradation remain significant obstacles to extended clinical use. Immune cell activation has been correlated to the acidity of breakdown products, yet methods to neutralize the pH have not significantly reduced adverse responses. Using a bioenergetic model, we observed delayed cellular changes that were not apparent in the short-term. Amorphous and semi-crystalline PLA degradation products, including monomeric L-lactic acid, mechanistically remodel metabolism in cells leading to a reactive immune microenvironment characterized by elevated proinflammatory cytokines. Selective inhibition of metabolic reprogramming and altered bioenergetics both reduce these undesirable high cytokine levels and stimulate anti-inflammatory signals. Our results present a new biocompatibility paradigm by identifying metabolism as a target for immunomodulation to increase tolerance to biomaterials, ensuring safe clinical application of PLA-based implants for soft- and hard-tissue regeneration, and advancing nanomedicine and drug delivery.

---

Polylactide (PLA) is the most widely utilized biopolymer^1^, with applications in nanotechnology, drug delivery and adult reconstructive surgery for tissue regeneration. However, after surgical implantation, PLA elicits adverse immune responses in up to 44% of human patients, often requiring further interventions^2,3^. In animals, a 66% incidence of excessive fibrosis with capsules from long-term inflammation which significantly limit implant-tissue integration has been reported^4^. PLA degrades by hydrolysis into D- or L-lactic acid, with semi-crystalline PLA degrading slower and tending to contain less D-content than amorphous PLA^1,5^. Adverse responses to PLA are exacerbated by mechanical loading and increasing implant size^6^, and occur after prolonged exposure to large amounts of PLA degradation products^2,7–9^. It is speculated that adverse responses are mediated by PLA degradation reducing pH in surrounding tissue^10^, the historical basis of which involved *Photobacterium phosphoreum*^11^. This bacterium expresses a luciferase whose reduced metabolic activity, measured by bioluminescence, can infer toxicity. In this study, breakdown products (extract) of PLA were obtained either in sterile water or Tris buffer; addition of acidic extract correlated with reduced luminescence. However, the study was not performed on mammalian cells, did not reflect the buffered in-vivo microenvironment or simulate prolonged exposure times to accumulated PLA degradation products. Establishing that a decrease in pH correlates with PLA degradation has informed the current strategy in regenerative medicine to neutralize acidic PLA degradation products both in-vitro and in-vivo using polyphosphazene^12^, calcium carbonate, sodium bicarbonate and calcium hydroxyapatite salts^10^, bioglass^13^ and composites containing alloys or hydroxides of magnesium^14,15^ despite reports of failures^16^. The lack of a clearly described mechanism of immune cell activation by PLA degradation remains a major obstacle in the safe application of large-PLA based implants in load-bearing applications as reflected by their paucity in FDA approvals^17^, and in soft tissue surgery where neutralizing ceramics cannot be applied^18^.

Metabolic reprogramming refers to significant changes in oxidative phosphorylation and glycolytic flux patterns and is a driver of fibrosis and bacterial lipopolysaccharide (LPS)-induced inflammation^19,20^. Here we set out to establish a molecular mechanism that directly links metabolic reprogramming to inflammation and fibrosis, consequent to cellular interactions with PLA degradation products. Foremost, we develop and validate a bioenergetic model of prolonged immune cell interaction with accumulated PLA degradation products. Only after prolonged exposure to amorphous or semi-crystalline PLA degradation products did macrophages and fibroblasts mechanistically undergo metabolic reprogramming and marked bioenergetic changes, with higher PLA crystallinity delaying onset. Using our model, we observed that PLA breakdown products markedly increase proinflammatory cytokine expression in primary macrophages through lactate signaling. Targeting different glycolytic steps using small molecule inhibitors modulated proinflammatory and stimulated anti-inflammatory cytokine expression by inhibiting metabolic reprogramming and altered bioenergetics in a dose-dependent manner. This process is highly specific and not cytotoxic to surrounding unaffected immune cells. Further, we demonstrate that use of the small molecule inhibitors imbedded in PLA implants substantiated our hypothesis of controlling the inflammatory response in-vivo. Our findings establish a new biocompatibility paradigm by identifying altered metabolism as a target for immunomodulation of PLA-based implants, fundamentally differing from previous strategies aimed at neutralizing PLA. Therefore, major advances in the use of PLA for human and veterinary applications are anticipated.

## Bioenergetic model for evaluating cellular responses to PLA degradation

To simulate in-vivo buffer conditions, breakdown products of PLA, generally referred to as extracts^21^, were generated in serum-containing DMEM medium and used after 12 days (d) of incubation in a shaker at 37 °C (Fig. 1a). This in-vitro degradation method was designed to mimic PLA degradation in-vivo, with agitation to accelerate PLA degradation relative to static methods^22^. Due to the buffering inherent in the serum-containing DMEM medium, there were no changes in pH over the 12 d extraction period for serum-containing control medium (pH = 8.0), amorphous PLA (pH = 8.2) and crystalline PLA (pH = 8.2) extracts used on cells. On the other hand, extraction in water for the same duration resulted in pH differences between control (pH = 8.2), amorphous PLA (pH = 7.5) and crystalline PLA (pH = 7.6) extracts.

**Figure 1.**
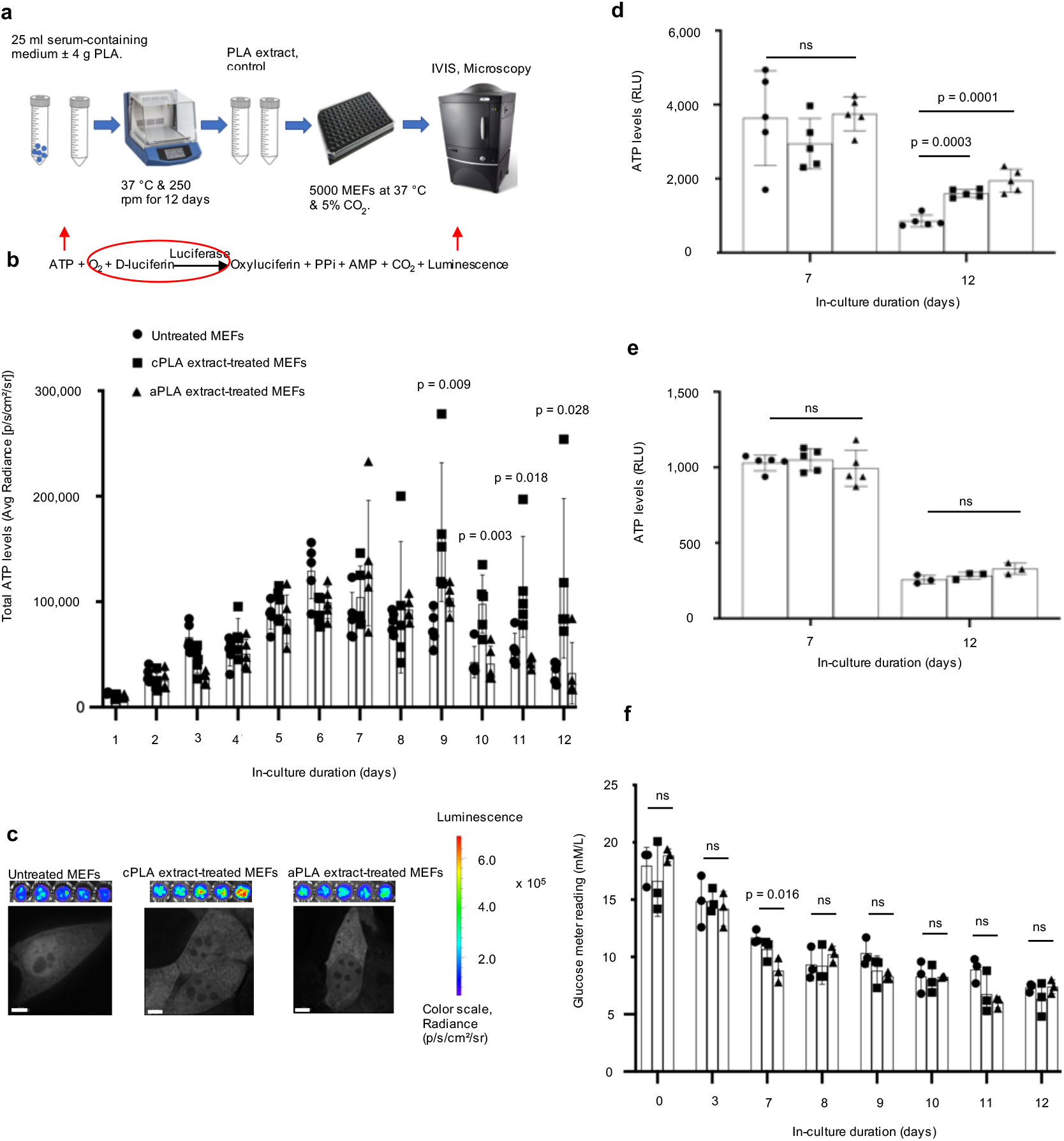
Bioenergetic (ATP) levels are elevated in mouse embryonic fibroblasts (MEFs) only after prolonged exposure to polylactide (PLA) degradation products (extract). **a**, Workflow showing our in-vitro bioenergetic model. **b**, Keeping luciferase, oxygen and D-luciferin levels constant (red circle) allows for changes in ATP (red arrow) to be measured by luminescence (red arrow). Using in-vivo imaging system (IVIS) and in comparison to controls, ATP levels in live cells are increased in blasticidin-eGFP-luciferase (BGL)-transfected MEFs after prolonged exposure to crystalline PLA (cPLA) degradation products. **c**, Representative microscopic (scale bars, 5 μM) and IVIS images show differential nucleoli number and luminescence, respectively. **d**, Measuring ATP in cell lysates of wild-type MEFs revealed that prolonged exposure to both amorphous PLA (aPLA) and cPLA results in elevated ATP levels. **e**, Addition of PLA does not affect the biochemical reaction by which ATP is measured. **f**, Between groups on the same day, glucose levels are similar in our in-vitro bioenergetic model. Not significant (ns), mean (SD), n = 5 (Fig. 1b, 1d and day 7 for 1e) or n = 3 (Fig. 1f and day 12 for 1e), one-way ANOVA followed by Tukey’s post-hoc test; 100 μl of control or PLA extract was used.

Together, studies in rodents, dogs and humans indicate that adverse immune responses occur after accumulation of PLA degradation products over several weeks or months^8,23–25^. To account for these extended exposure times in our model, we cultured immune cells in PLA extract for 12 d, and this required initiating our cultures with small numbers of cells per well in both control and treatment groups to prevent overgrowth of the cultures. Mouse embryonic fibroblasts (NIH 3T3 cells) were stably transfected with a Sleeping Beauty transposon plasmid (pLuBIG) having a bidirectional promoter driving a modified firefly luciferase gene (fLuc) and a fusion gene encoding a Blasticidin-resistance marker (BsdR) linked to eGFP (BGL)^26^. Seeding the same cell numbers across control and treatment groups resulted in constant levels of luciferase and we exposed cells to equal levels of D-luciferin and oxygen in all assays. In this manner, ATP was rate-limiting and changes in ATP were measured by bioluminescence using in-vivo imaging system (IVIS; Fig. 1b). Use of bioluminescence as an indicator of ATP levels was inexpensive, rapid (on the order of seconds) and allowed for high throughput temporal bioenergetic analysis in live cells. Additionally, in our model, each well of a 96-well plate had a total of 200 *μ*l of medium, of which 100 *μ*l was freshly prepared. The additional 100 *μ*l for control wells was medium that had been in the shaker at 37 °C for 12 d to account for potential nutrient degradation that could confound results. Similarly, the additional 100 *μ*l for treatment wells was medium in which PLA had been degraded under the same conditions.

Dose-bioenergetic response of amorphous and crystalline PLA extracts revealed altered ATP levels for all tested doses (Supplementary Fig. 1a). Therefore, we selected 100 or 150 *μ*l of extract, as indicated in figure legends, to mimic accumulation of voluminous PLA breakdown products^2–7^.

Highly crystalline and amorphous PLA samples were selected for their high molecular weights and represent a range of physicochemical properties (crystallinity, stereochemistry, degradation period) which constitute important considerations in selecting PLA for hard and soft tissue engineering^8,10,25^. Before using these PLA materials, we authenticated their physicochemical and thermal properties (Supplementary Table 1). Lastly, we used the non-transformed, immortalized NIH 3T3 fibroblast cell line that typifies primary fibroblasts, as well as primary bone-marrow derived macrophages, both of which are key cellular mediators of prolonged inflammation and excessive fibrosis that occur in response to PLA degradation^12,23^.

## Bioenergetics is altered in immune cells after exposure to PLA degradation products

Unlike in the short-term (days 0-5), prolonged (days 6-12) exposure of fibroblasts to either amorphous or crystalline PLA increased ATP levels in live cells (Fig. 1b-c). Upon high resolution z-stack imaging, there were apparent changes in nucleoli number (Fig. 1c) after prolonged exposure to either amorphous or crystalline PLA extract, which could represent a stress response^27^. To exclude the possibility that changing luciferase expression (by transcription or translation) was responsible for observed bioenergetic changes, we lysed wild-type cells after exposure to PLA extract and added controlled amounts of luciferase and D-luciferin in the standard ATP assay. Moreover, measuring ATP levels in live cells by IVIS is constrained by parameters inherent to live cells. Lysed cells allow for the measurement of ATP from all organellar compartments, and is not constrained by D-luciferin uptake, revealing more information than measurements in live cells. By day 12, there was a 1.9- and 2.3-fold increase in ATP levels among cells exposed to crystalline and amorphous PLA extract, respectively (Fig. 1d). To exclude the possibility that PLA extracts affect the biochemical reaction (Fig. 1b) underlying bioenergetic measurements, fibroblasts were cultured for different time points. Thereafter, lysed fibroblasts were exposed to D-luciferin, luciferase and control or PLA extract at the same time. No difference in ATP levels was observed, confirming that treatment with PLA extract did not affect this biochemical reaction (Fig. 1e).

Declining ATP levels from day 0 to day 12 is likely due to changing glucose levels^28^. To determine whether glucose levels changed between groups on the same day because of the extended exposure times in our model, glucose meter readings were optimized in mammalian cell culture medium (Supplementary Fig. 1b). Glucose levels were similar between groups on each day (Fig. 1f). On day 7, when untreated groups had higher glucose levels (Fig. 1f), corresponding bioenergetic measurement revealed that PLA extract-treated fibroblasts had higher ATP levels (Supplementary Fig. 1c), excluding changing glucose levels as a confounding factor in our bioenergetic model. Because NIH 3T3 cells are normal immortalized fibroblasts, changing cell number from proliferation could account for bioenergetic changes. To exclude this, we optimized the crystal violet assay for cell number measurement^29^ in fibroblasts (Supplementary Fig. 2a). Next, we isolated mouse primary bone marrow-derived macrophages (BMDMs) which, unlike NIH 3T3 cells, do not proliferate^30^.

Both ATP^31^ and ADP^32^ metabolism and ratios are crucial in inflammatory conditions. In BMDMs and consistent with our observations in fibroblasts, we observed marked increases in ATP and ADP levels (Fig. 2a, b) or ATP/ADP ratios (Fig. 2c) which were not due to changing glucose levels (Fig. 2d). After optimizing the crystal violet assay for macrophages (Supplementary Fig. 2b), overall, cell numbers could not account for observed bioenergetic changes (Fig. 2e). Furthermore, fibroblast numbers were similar for cultures that were untreated or exposed to PLA extracts (Fig. 2f), excluding changing cell number as a confounder in our model.

**Figure 2.**
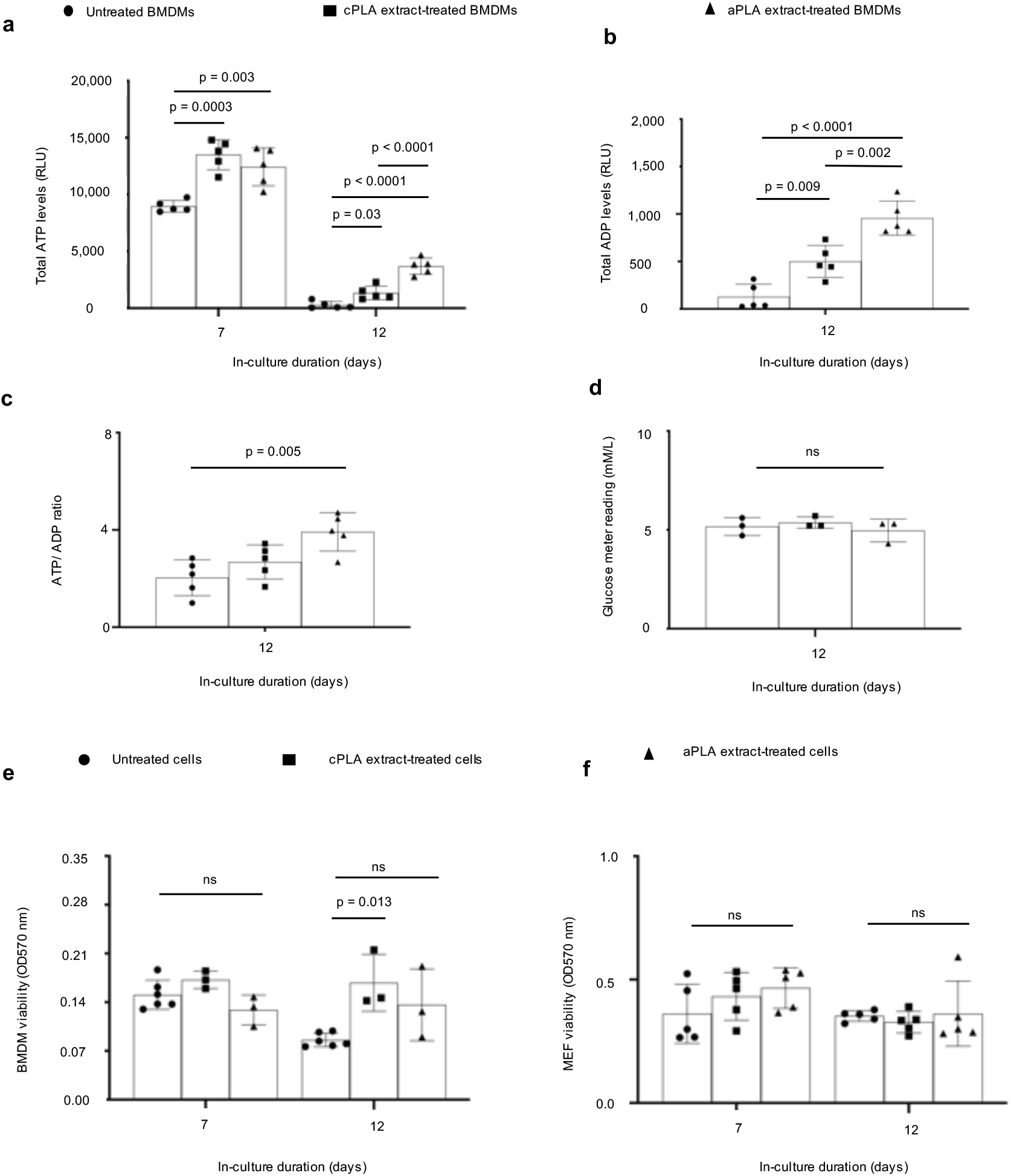
Bioenergetics is increased in primary bone marrow-derived macrophages (BMDMs) after prolonged exposure to polylactide (PLA) degradation products (extract). **a**, ATP levels **b**, ADP levels **c**, and ATP/ADP ratios are increased in BMDMs after prolonged exposure to amorphous PLA (aPLA) or crystalline PLA (cPLA) degradation products (extracts) in comparison to controls. **d**, Glucose levels between groups on day 12 are similar. **e-f**, Cell number between groups are similar for BMDMs (**e**) and MEFs (**f**). Not significant (ns), mean (SD), n = 5 (Fig. 2a, b, c, f), n = 3 (Fig. 2d), n = 3-6 (Fig. 2e), one-way ANOVA followed by Tukey’s post-hoc test; 100 μl of control or PLA extract was used.

## Exposure of macrophages to PLA breakdown products selectively results in metabolic reprogramming

To determine the metabolic pathways responsible for the bioenergetic changes we had observed, Seahorse assays were used to measure oxygen consumption rate (OCR), extracellular acidification rate (ECAR) and lactate-linked proton efflux rate (PER) in a customized medium (pH 7.4); this technique has not been previously used to examine PLA-induced adverse responses. PLA extract was removed and washed off the cells prior to running the Seahorse assay at a pH of 7.4. Seahorse assays measure ECAR as an index of glycolytic flux, OCR as an index of oxidative phosphorylation and PER as an index of monocarboxylate transporter function^33^ in live cells; and are used to assess for metabolic reprogramming^34–36^. Primary BMDMs exposed to amorphous PLA extract were metabolically altered, showing a 2-fold increase in oxidative phosphorylation (OCR; Fig. 3a), 3.5-fold increase in glycolytic flux (ECAR; Fig. 3b) and 3.5-fold increase in monocarboxylate transporter activity (PER; Fig. 3c) in comparison to untreated BMDMs. Similar amounts (100 μl) of crystalline PLA extract resulted in a 1.6-fold increase in OCR (Fig. 3d) but no change in ECAR (Fig. 3e) or PER (Fig. 3f). However, higher amounts (150 μl) of crystalline PLA extract resulted in 3.2-, 3.8-, and 3.8-fold increases in OCR, ECAR and PER, respectively (Supplementary Fig. 3a-c) compared to controls, suggesting that greater volume of PLA extract is required for reprogramming using crystalline than amorphous PLA.

**Figure 3.**
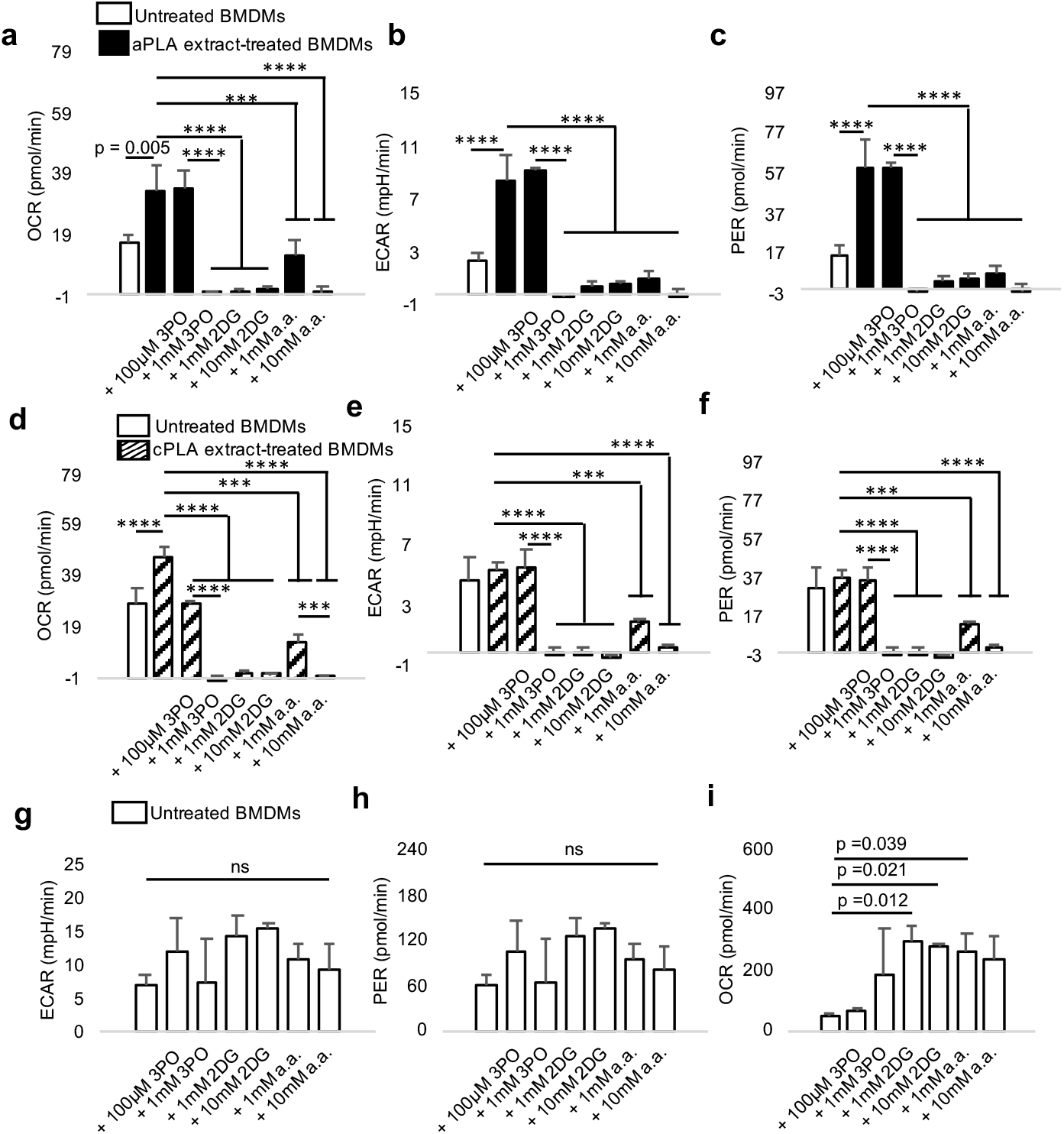
Functional metabolic indices are altered in primary bone marrow-derived macrophages (BMDMs) after prolonged exposure to polylactide (PLA) degradation products (extract), and can be modulated by glycolytic inhibitors. **a-c**, Following exposure to amorphous PLA (aPLA) extract, oxygen consumption rate (OCR) (**a**), extracellular acidification rate (ECAR) (**b**) and proton efflux rate (PER) (**c**) are increased relative to controls, and this abnormal increase can be dose-dependently controlled by various small molecule inhibitors. **d-f**, OCR (**d**) and not ECAR (**e**) and PER (**f**) are increased relative to controls in groups exposed to crystalline PLA (cPLA) extract, and functional metabolic indices can be controlled by pharmacologic inhibitors of glycolysis. **g-h**, ECAR (**g**) and PER (**h**) are not affected by glycolytic inhibitors in untreated BMDMs. **i**, Compensatory increase in OCR occurs in untreated BMDMs after treatment with some inhibitors. Not significant (ns), ***p<0.001, ****p<0.0001, mean (SD), n = 3, one-way ANOVA followed by Tukey’s post-hoc test; 3-(3-pyridinyl)-1-(4-pyridinyl)-2-propen-1-one (3PO), 2-deoxyglucose (2DG) and aminooxyacetic acid (a.a.); 100 μl of control or PLA extract was used for 7 days.

Next, we targeted different steps in the glycolytic pathway using three small molecule inhibitors: 3-(3-pyridinyl)-1-(4-pyridinyl)-2-propen-1-one (3PO), 2-deoxyglucose (2DG) and aminooxyacetic acid (a.a.). Whereas 3PO specifically inhibits 6-phosphofructo-2-kinase which is the rate limiting glycolytic enzyme^37^, 2DG inhibits hexokinase, the first enzyme in glycolysis^36^, and aminooxyacetic acid prevents uptake of glycolytic substrates^38^. In a dose-dependent manner, 3PO, 2DG and a.a. inhibited metabolic reprogramming following exposure to amorphous PLA (Fig. 3a-c) or crystalline PLA extract (Fig. 3-f), but not in untreated BMDMs (Fig. 3g-i).

This demonstrates cellular uptake of 3PO, 2DG and a.a., yet with selective pharmacologic effects. Notably and under the same experimental conditions, cell viability was not reduced in untreated BMDMs after exposure to glycolytic inhibitors (Supplementary Fig. 2c), demonstrating the absence of cytotoxicity^29^. However, when BMDMs were treated with amorphous or crystalline PLA extract, where metabolism was abnormally remodeled, 3PO, 2DG and a.a. mildly, but selectively, reduced cell viability (Supplementary Fig. 2d). Therefore, pharmacologically targeting altered metabolism in primary BMDMs following exposure to PLA extract is highly specific with limited toxicity to immune cells that have normal metabolic profiles.

## Fibroblasts are glycolytically reprogrammed after exposure to PLA breakdown products

After prolonged exposure of fibroblasts to amorphous and crystalline PLA extracts, glycolytic flux (ECAR; Fig. 4a-b) is increased by 1.6- and 1.7-fold, respectively. Furthermore, monocarboxylate transporter function is increased in amorphous or crystalline PLA extract-treated fibroblasts by 1.6- and 1.5-fold, respectively (Fig. 4c-d). However, oxidative phosphorylation remains similar between untreated fibroblasts and cells exposed to amorphous or crystalline PLA extracts (OCR; Supplementary Fig. 4a-b). Remarkably, increased bioenergetic (ATP) levels in amorphous or crystalline PLA extract-treated fibroblasts are inhibited by 3PO, 2DG and a.a. in a temporal and dose-dependent manner (Fig. 4e; Supplementary Fig. 4c).

**Figure 4.**
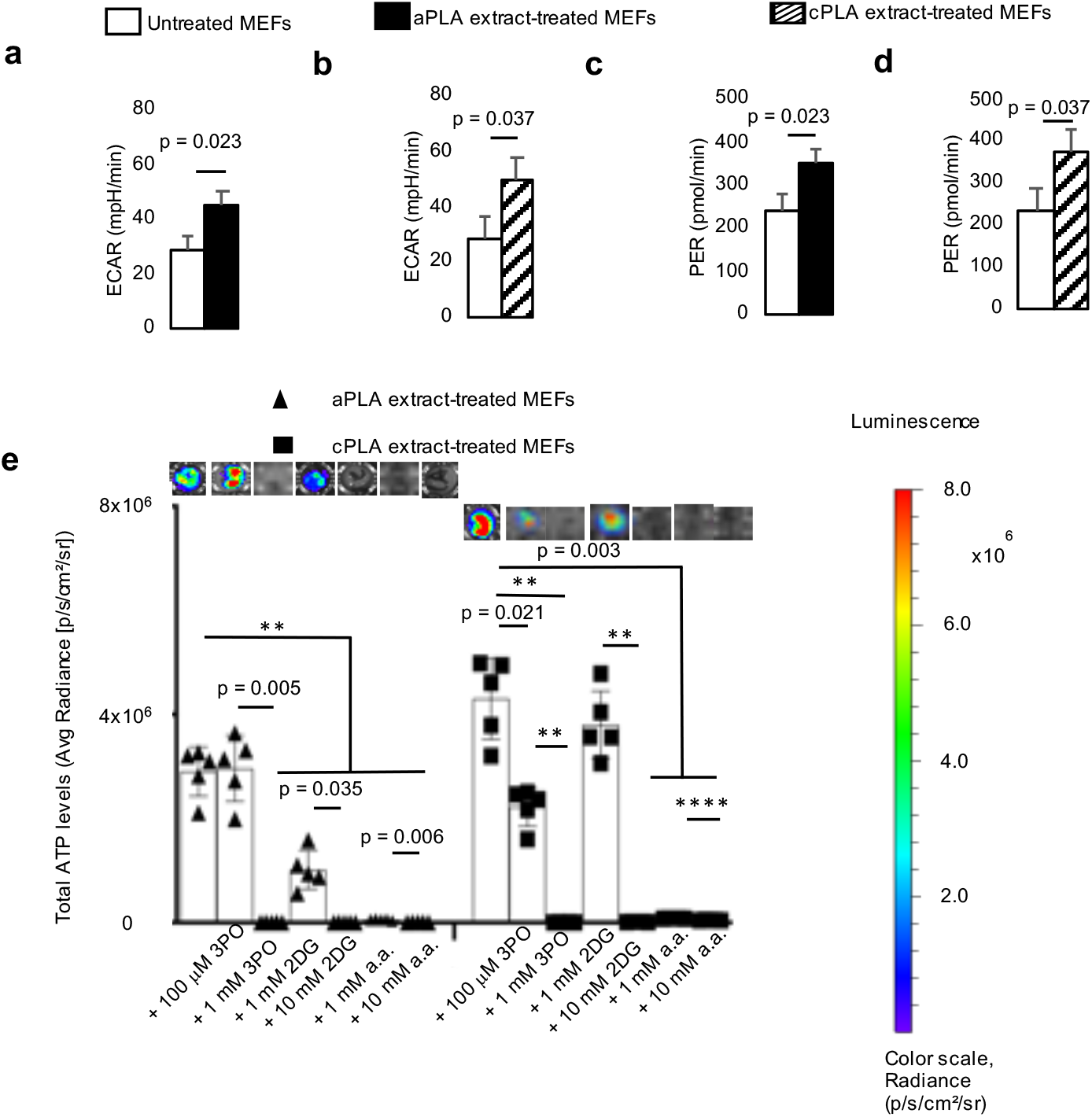
Functional metabolism is altered in mouse embryonic fibroblasts (MEFs) after exposure to polylactide (PLA) degradation products (extract). **a-b**, Following exposure to amorphous PLA (aPLA; **a**) or crystalline PLA (cPLA; **b**) extracts, extracellular acidification rate (ECAR) is increased. **c-d**, Proton efflux rate (PER) is elevated in MEFs after exposure to aPLA (**c**) or cPLA (**d**) extract. **e**, Bioenergetic levels in MEFs exposed to aPLA or cPLA extracts are decreased in a dose-dependent manner by 3-(3-pyridinyl)-1-(4-pyridinyl)-2-propen-1-one (3PO), 2-deoxyglucose (2DG) and aminooxyacetic acid (a.a.; representative wells are shown). ** p = 0.002, **** p < 0.0001, mean (SD), n = 3 (Fig. 4a, b, c, d), n = 5 (Fig. 4e), two-tailed unpaired t-test or Brown-Forsythe and Welch ANOVA followed by Dunnett’s T3 multiple comparisons test; 100 μl of control or PLA extract was used for 7 days.

## Short- and long-term exposure to L-lactic acid alters bioenergetics and results in metabolic reprogramming

As previously reported for short-term hydrolytic degradation of PLA^8^, there was no reduction in mass of PLA after our 12 d extraction, but there were detectable changes in molecular weight (Supplementary Table 2). Using the standard D/L-lactic acid enzyme-based determination assays could not effectively measure levels in serum-containing medium. However, in milliQ water and relative to controls, we observed a 7.8- and 5.2-fold increase in L-lactic acid in amorphous and crystalline PLA extracts, respectively, although these increments were not significant (Supplementary Table 3; Supplementary Fig. 5a-b). Similarly, we observed a 2.7- and 2.8-fold increase in D-lactic acid in amorphous and crystalline PLA extracts, respectively (Supplementary Table 3). Therefore, we exposed BMDMs to various doses of L-lactic acid, ranging from 2.5- to 15-fold higher levels in comparison to untreated cells. In addition, we measured corresponding pH levels: Untreated medium (pH = 8.01), 2.5 mM (pH = 7.47), 5 mM (pH = 7.19), 10mM (pH = 6.84) and 15mM (pH = 6.65) L-lactic acid-containing DMEM medium.

We observed that bioenergetic levels are altered in the short-term (day 3; Fig. 5a) for all doses of L-lactic acid treatment, resulting in a 1.5 to 1.6-fold increase in ATP levels. After prolonged (day 7) exposure to L-lactic acid and even when bioenergetic alterations were not apparent, glycolytic flux (ECAR; Fig. 5b), monocarboxylate transporter function (PER; Fig. 5c) and oxidative phosphorylation (OCR; Fig. 5d) were increased by 2.8-, 2.8- and 2.3-fold, mechanistically reproducing observations made with amorphous and crystalline PLA extracts in our bioenergetic model. Moreover, these changes were not dependent on alterations in cell number (Supplementary Fig. 5c). Of note, highly acidic groups (10-15 mM L-lactic acid) did not result in reduction in viability of primary macrophages either at day 7 or 12, relative to controls (Supplementary Fig. 5c).

**Figure 5.**
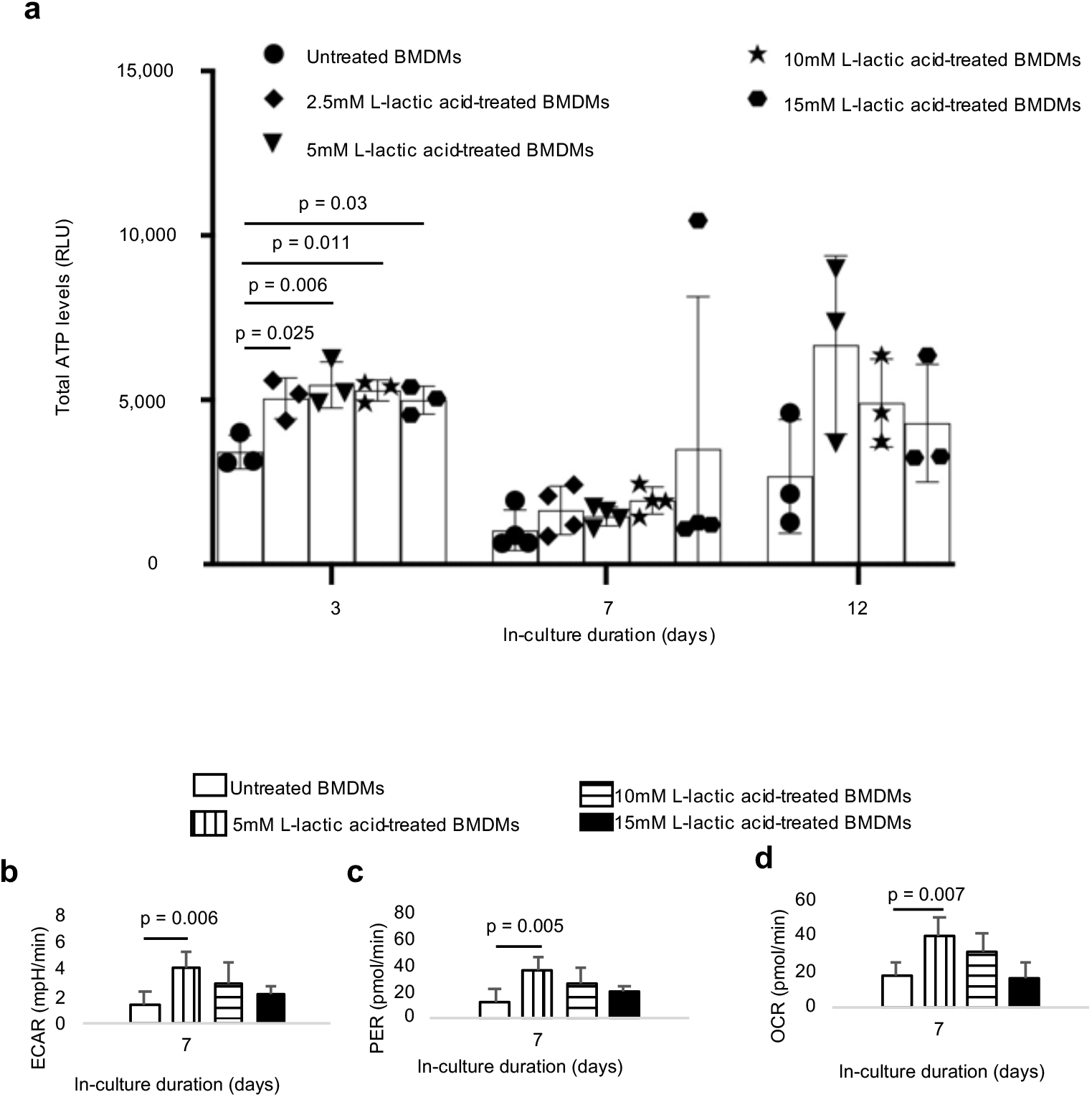
Treatment of primary bone marrow-derived macrophages (BMDMs) with L-lactic acid altered bioenergetic (ATP) levels and functional metabolism. **a**, Treatment with different doses of monomeric L-lactic acid resulted in changes in ATP levels. **b-d**, Following exposure to L-lactic acid extracellular acidification rate (ECAR, **b**), proton efflux rate (PER, **c**) and oxygen consumption rate (OCR, **d**) are increased. One-way ANOVA followed by Tukey’s post-hoc test, mean (SD), n = 3-4 (Fig. 5a), n = 5 (Fig. 5b, c, d).

## Glycolytic inhibition modulates proinflammatory and stimulates anti-inflammatory cytokine expression

To determine whether glycolytic inhibition affects proinflammatory (IL-6, MCP-1, TNF-α, IL-1 β and IFN-l) and anti-inflammatory (IL-4, IL-10, and 1L-13) protein expression, we used a magnetic bead-based chemokine and cytokine assay^39^. We observed that prolonged exposure of primary macrophages to amorphous and crystalline PLA extracts resulted in 228- and 319-fold increases, respectively, in IL-6 protein expression (Fig. 6a) compared to untreated macrophages. We confirmed this observation by ELISA (Supplementary Fig. 6a). Similarly, exposure of macrophages to lactic acid resulted in elevated IL-6 protein expression by 2.3-fold (Supplementary Fig. 6a). Amorphous PLA extracts increased MCP-1 (Fig. 6b), TNF-α (Fig. 6c) and IL-1β (Fig 6d) levels by 1.2-fold, 21-fold, and 567-fold, respectively. Likewise, crystalline PLA extracts increased MCP-1 (Fig. 6b), TNF-α (Fig. 6c) and IL-1β (Fig 6d) levels by 4.7-fold, 27-fold, and 1,378-fold, respectively. Abnormally increased levels of IL-6, MCP-1, TNF-α and IL-1β were modulated by addition of 3PO, 2DG or a.a. (Fig. 6a-d). Increased MCP-1 levels in macrophages also occurred after exposure to L-lactic acid (Supplementary Fig. 6b). Levels of IFN-γ and IL-13 were unchanged by PLA extract (data not shown) but exposure to amorphous PLA extract decreased IL-4 protein levels by 3-fold (Fig. 6e) relative to untreated macrophages. Remarkably, with the exception of 3PO, IL-10 expression was either unchanged (crystalline PLA) or increased by 3.4-fold (amorphous PLA) upon addition of a.a. (Fig. 6f) relative to macrophages exposed to PLA extract.

**Figure 6.**
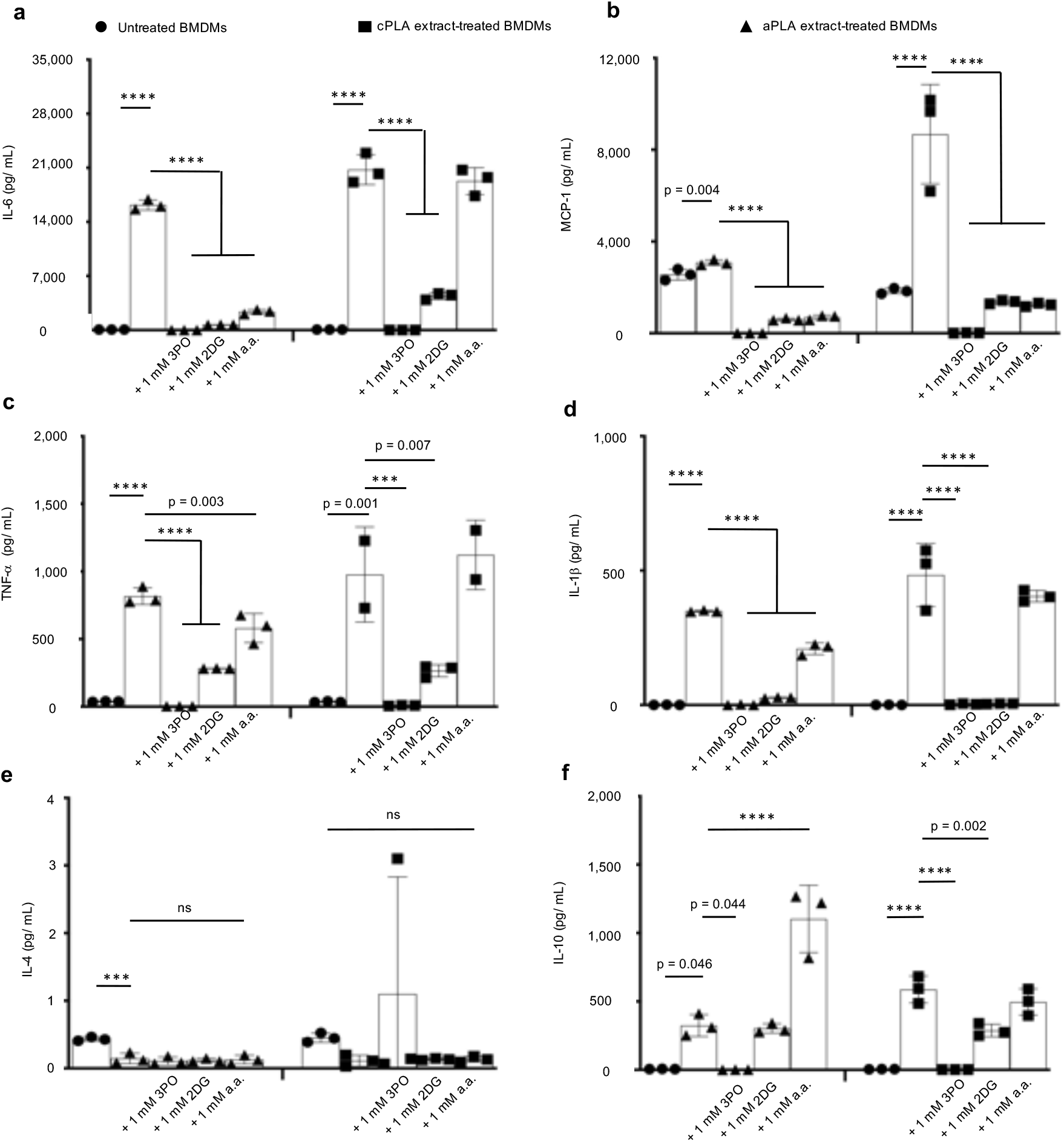
In macrophages exposed to PLA breakdown products, glycolytic inhibitors modulate elevated proinflammatory cytokine expression and stimulate or do not reduce anti-inflammatory cytokine levels. **a-d**, Following exposure to amorphous PLA (aPLA) or crystalline PLA (cPLA) extract, primary bone marrow-derived macrophages (BMDMs) express elevated levels of IL-6 (**a**), MCP-1 (**b**), TNF-α (**c**) and IL-1 β (**d**) in comparison to untreated BMDMs, and these elevated proinflammatory cytokine levels can be modulated by various small molecule inhibitors of glycolysis. **e,** Addition of glycolytic inhibitors to PLA does not reduce IL-4 expression. **f**, Expression of IL-10 is increased by inhibiting glycolysis using aminooxyacetic acid (a.a.) in amorphous PLA. Not significant (ns), ***p<0.001, ****p<0.0001, mean (SD), n = 3 in all except the cPLA group in TNF-α (Fig. 6c) where n=2-3, one-way ANOVA followed by Tukey’s post-hoc test; 3-(3-pyridinyl)-1-(4-pyridinyl)-2-propen-1-one (3PO), 2-deoxyglucose (2DG); 100 μl of aPLA or 150 μl of cPLA extract with corresponding controls were used on day 7.

## Increased radiolabeled glucose uptake occurs in the PLA microenvironment and drives inflammation, in-vivo

Taken together, our in-vitro data suggest that metabolic changes drive inflammation arising from PLA degradation. To test this hypothesis in-vivo, we incorporated 2DG into amorphous PLA (aPLA) by melt-blending at 190 °C and compared to aPLA which had been subjected to similar melt-blending conditions (called reprocessed aPLA). Following melt-blending, extruded (sterile) filaments (1.75 mm diameter, 1 mm long) were subcutaneously implanted on the back (dorsum) of mice. Sham controls underwent similar surgical exposures but were not implanted with any materials. After 6 weeks, mice were injected with F-18 fluorodeoxyglucose (FDG) and euthanized; using FDG allows for evaluation of metabolic reprogramming and inflammation^40,41^. Thereafter, circular biopsies (12 mm in diameter) of full thickness skin containing implants were assayed for radioactivity using a gamma counter. Compared to sham controls, skin containing reprocessed aPLA implants had 1.35-fold increase in FDG uptake, which was abolished in skin containing aPLA+2DG implants (Fig. 7a).

**Figure 7.**
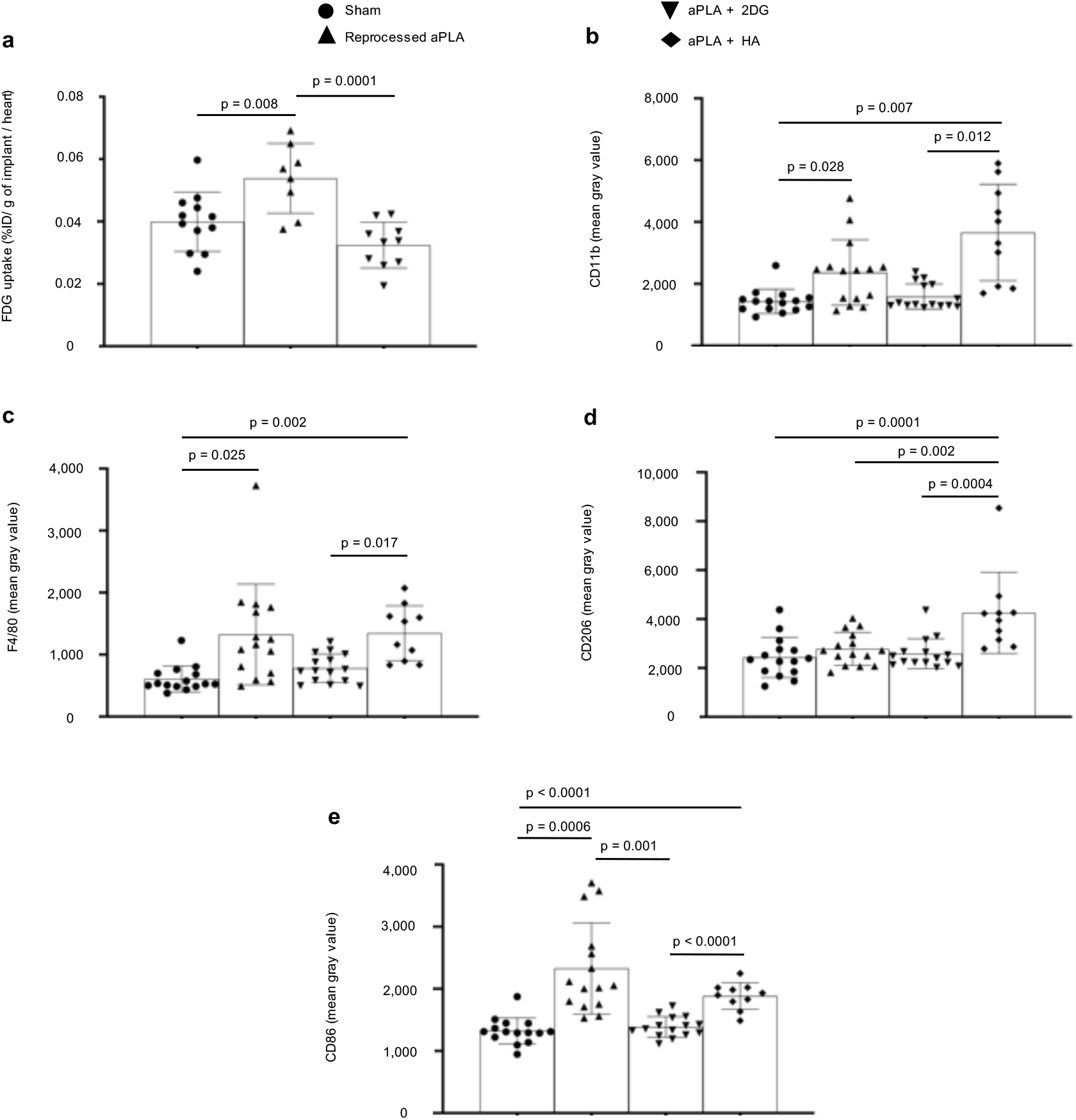
Increased radiolabeled glucose uptake occurs in the polylactide (PLA) microenvironment and drives inflammation in-vivo. **a**, When normalized to heart values, percent injected dose per gram (% ID/ g) of biopsied tissues surrounding amorphous PLA (aPLA) implants show higher F-18 fluorodeoxyglucose (FDG) uptake compared to sham controls; increased FDG uptake is reduced by incorporation of 2-deoxyglucose (2DG). **b,c**, Compared to sham controls, mean fluorescence intensity of CD11b (**b**) or F4/80 (**c**) is increased following surgical implantation of aPLA or a combination of aPLA and hydroxyapatite (HA), but not a combination of aPLA and 2DG. **d**, Compared to other groups, CD206 mean fluorescence intensity is increased in aPLA + HA. **e**, Compared to sham controls, CD86 mean fluorescence intensity is increased following implantation of aPLA elevated CD86 is decreased by incorporating 2DG but not HA. Mean (SD); Fig. 1a, sham (n = 12), aPLA (n = 8), aPLA + 2DG (n = 10); Fig 1b-e, sham (n = 15), aPLA (n = 15), aPLA + 2DG (n = 15), aPLA + HA (n = 10); one-way ANOVA followed by Tukey’s post-hoc test or Brown-Forsythe and Welch ANOVA followed by Dunnett’s T3 multiple comparison test; refer to methods section (In-vivo studies, tissue processing and analyses) for more information on n.

Next, we sought to determine the effect of glycolytic inhibition on recruitment and activation states of macrophages and fibroblasts. To compare the effects of glycolytic inhibition to neutralization techniques, we included a group where hydroxyapatite (HA) was incorporated in aPLA^10,42^. Hematoxytin and eosin staining revealed the presence of inflammatory infiltrates in the implant microenvironment (reprocessed aPLA, aPLA+2DG, aPLA+HA) compared to sham controls, suggesting persistent inflammation (Supplementary Fig. 7).

Chronic inflammation to PLA is principally driven by recruited macrophages^12,23^. Therefore, we stained for CD11b and F4/80, established macrophage markers. Compared to sham controls, aPLA resulted in a 1.7- and 2.2-fold increase in CD11b and F4/80 intensities, respectively (Fig. 7b-c; Supplementary Fig. 8). Unlike aPLA+2DG, aPLA+HA increased CD11b and F4/80 intensities by 2.6-fold and 2.2-fold, respectively, when compared to sham controls (Fig. 7b-c). Of note, there was no significant difference in CD11b and F4/80 intensities between aPLA and aPLA+2DG (Fig. 7b-c), suggesting similar levels of macrophage recruitment. Furthermore, aPLA+2DG revealed 2.3-fold and 1.7-fold less CD11b and F4/80 intensities, respectively, compared to aPLA+HA (Fig. 7b-c). To determine the activation states of recruited macrophages, we stained for CD206 and CD86, anti-inflammatory and proinflammatory macrophage markers, respectively^43^. Relative to other groups, only aPLA+HA increased CD206 intensity (Fig. 7d; Supplementary Fig. 8), consistent with the bioactivity of HA^44^. We observed a 1.8-fold increase in CD86 intensity with reprocessed aPLA compared to sham controls, consistent with the proinflammatory effects of aPLA (Fig. 7e; Supplementary Fig. 8). Compared to reprocessed aPLA, aPLA+2DG and not aPLA+HA decreased CD86 intensity (Fig. 7e). In fact, there was a 1.4-fold decrease in CD86 intensity in aPLA+2DG compared to aPLA+HA (Fig. 7e).

Fibroblasts are a key cellular player of excessive fibrosis around PLA implants^12,23^, and their activation in myofibroblast phenotype is marked by a-SMA and TGF-β expression^45^. We observed a 1.4-fold increase in a-SMA intensity with reprocessed PLA compared to sham controls, which was decreased in the aPLA+2DG, but not aPLA+HA group (Fig. 8a; supplementary Fig. Supplementary Fig. 9). With TGF-β intensity, aPLA+HA was elevated relative to other groups (Fig. 8b; Supplementary Fig. 9). Compared to aPLA+HA, aPLA+2DG revealed 1.4-fold and 1.8-fold decrease in a-SMA and TGF-β intensities, respectively (Fig. 8b).

**Figure 8.**
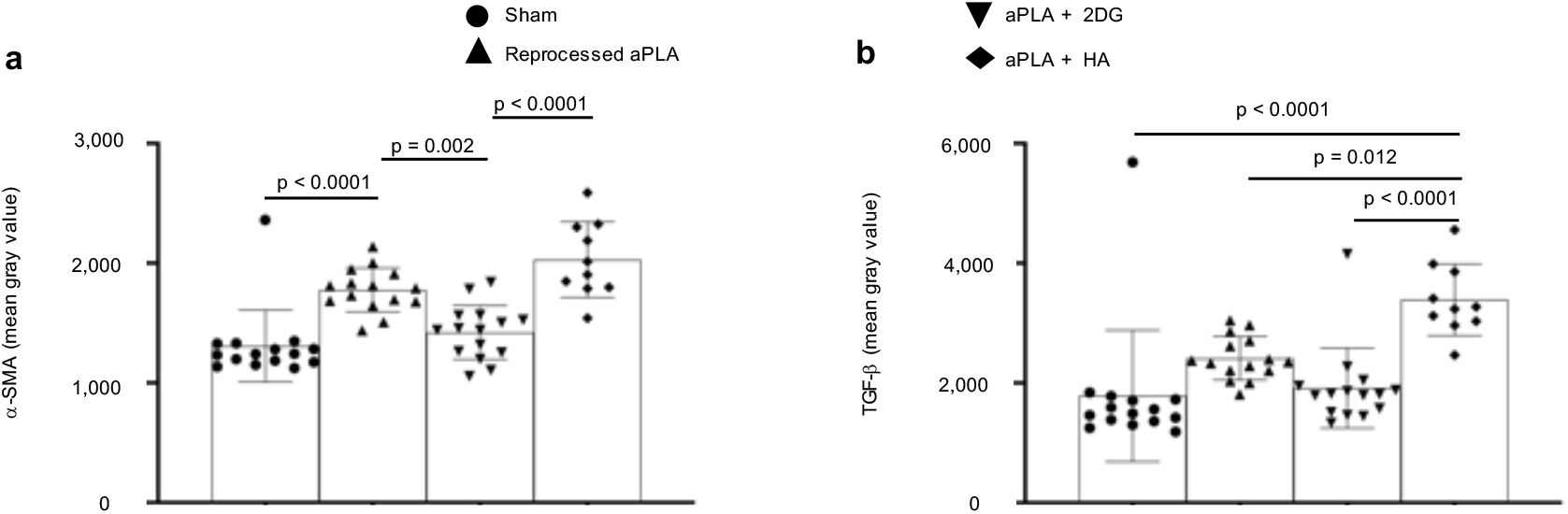
Activation of fibroblasts in the polylactide (PLA) microenvironment is regulated by immunometabolism. **a**, Compared to sham controls, mean fluorescence intensity of alpha-smooth muscle actin (α-SMA) is increased following surgical implantation of amorphous PLA (aPLA) or a combination of aPLA and hydroxyapatite (HA), but not a combination of aPLA and 2-deoxyglucose (2DG). **b**, Compared to other groups, mean fluorescence intensity of transforming growth factor-beta (TGF-β) is increased in aPLA + HA. Mean (SD); sham (n = 15), aPLA (n = 15), aPLA + 2DG (n = 15), aPLA + HA (n = 10); one-way ANOVA followed by Tukey’s post-hoc test; refer to methods section (In-vivo studies, tissue processing and analyses) for more information on n.

## Discussion

We describe a bioenergetic model of immune cell activation to PLA degradation, revealing that altered bioenergetics and metabolic reprogramming underlie adverse responses, including persistent inflammation and excessive fibrosis, to PLA breakdown products. For decades, the hypothesis in regenerative medicine has been that acidity drives immune cell activation to PLA degradation^10^. However, this observation was founded on correlation and not causation^11,21^. Consequently, methods based on neutralizing acidity have been inadequate in controlling adverse responses to PLA degradation^7,16^.

Importantly, our in-vitro model extends the short time periods that have been previously studied^46^. By adapting our bioenergetic model for high throughput analysis, we observed delayed immune cell changes not apparent in the short-term. In patients, PLA slowly degrades into oligomers and monomers of lactic acid. Ultimately, due to bulk degradation, PLA breakdown exceeds immune cellular clearance, resulting in accumulation of oligomers and monomers of lactic acid^25^. We illustrate that only after prolonged exposure to PLA degradation products do fibroblasts and macrophages become activated. Mechanistically, PLA degradation not only alters bioenergetic homeostasis in immune cells, it results in metabolic reprogramming. We identified PLA degradation products to include monomeric L-lactic acid and reproduced bioenergetic alterations and metabolic reprogramming using monomeric L-lactic acid. Although not shown, we also identified a spectrum of different oligomeric lactic acid in extracts by electrospray ionization-mass spectrometry as previously reported^47^. Following prolonged exposure of macrophages to PLA degradation products, metabolic reprogramming is characterized by concomitantly elevated oxidative phosphorylation and glycolysis, resulting in increased IL-6, MCP-1, TNF-α and IL-1β protein expression, potent proinflammatory cytokines. Increased glycolysis, a fundamental proinflammatory metabolic phenotype, is likely mediated by HIF-1a^36^. Human fibroblasts in lactate-enriched medium stabilize HIF-1a resulting in increased glycolysis^48^ which underlies activation of fibroblasts in several profibrotic pathologies^19^. Similarly, increased oxidative phosphorylation is required for macrophages to function as antigen presenting cells as part of inflammation^49^ or its resolution^20^.

Inhibiting different steps in the glycolytic pathway produced similar effects, decreasing proinflammatory cytokine expression by modulating metabolic reprogramming and altered bioenergetics. Unlike bacterial LPS-mediated glycolytic reprogramming that is uniquely dependent on IL-1β^36^, PLA degradation products additionally affect IL-6, MCP-1 and TNF-α. Of note, modulating proinflammatory cytokine expression using aminooxyacetic acid stimulated IL-10 protein expression, an anti-inflammatory cytokine^34^. Collectively, these findings are important for at least four reasons. First, it explains the “Oppenheimer phenomenon”, where long-term PLA implantation results in neoplasia in some humans and up to 80% of rodents^6^ since IL-6 directly links persistent inflammation from PLA to cellular transformation^50^. Second, stimulating IL-10 is critical to tissue repair by driving wound resolution and angiogenesis^51^. In fact, IL-10 is a key immunomodulatory cytokine secreted by mesenchymal stem cells^52^, and is crucial in macrophage-stem cell crosstalk^53,54^ for tissue engineering. Third, macrophages that have normal metabolism are unaffected by the small molecule inhibitors studied. In fact, cytotoxicity is selective for macrophages having altered metabolism, following exposure to PLA degradation products, making this technique particularly desirable. Fourth, it provides a basis to study lactate signaling in tumor initiation, with the potential to stop neoplastic initiation.

In cell culture medium used in our studies, serum and high bicarbonate salts buffered the pH of PLA degradation products, excluding pH as a confounder of observed metabolic cellular changes. Furthermore, using monomeric L-lactic acid at various concentrations that simulated neutralized and acidic PLA degradation products, we similarly observed bioenergetic alterations, excluding pH as a confounder. Lastly, using aminooxyacetic acid to modulate some adverse responses to PLA degradation products, suggests that acidity is not solely the driver of immune cellular activation to PLA.

Our findings using sterile implants present a perspective different than what is observed with bacterial endotoxin (LPS). Within 1h of exposure to very low endotoxin concentrations, significant metabolic changes characterized by increased glycolysis and decreased oxidative phosphorylation occurs^35^; with PLA degradation products, we only observed changes after several days of exposure, with distinct metabolism. Importantly, LPS decreases ATP levels^35,55^; in contrast, PLA degradation products (including monomeric L-lactic acid) increase ATP levels as shown in this study. Lastly, unlike PLA degradation products, LPS-mediated glycolytic reprogramming is reliant on IL-1β signaling^36^.

Lactate is a signaling molecule in immunity^56^ and cancer progression^57–59^. Its role when combined with LPS is conflicting, with reports of proinflammatory and anti-inflammatory effects^60,61^. However, a stand-alone ability of lactate to activate immune cells is novel, as prior inflammatory and cancer models did not simulate prolonged exposure times, a critical feature of the cancer and immune microenvironments.

Amorphous PLA which undergoes faster hydrolytic degradation than crystalline PLA results in quicker onset of metabolic reprogramming. Nonetheless, crystalline PLA does eventually result in metabolic remodeling and altered bioenergetics. Furthermore, our data implicate monocarboxylate transporters which mediate the bi-directional flux of lactate across cell membranes^33,60^.

Glucose is the first substrate in glycolysis. As such, radiolabeled glucose (FDG) uptake is often used to measure glycolytic dependence, in-vivo, such as in some cancers or inflammatory disorders where enhanced glycolysis is pivotal to disease progression^62^. We observed increased glycolytic dependence in the PLA inflammatory microenvironment using sterile amorphous PLA, which was abrogated by 2DG, one of the glycolytic inhibitors applied in our in-vitro studies. Unsurprisingly, after surgical resection of colorectal and cervical tumors in human patients, chronic, sterile inflammation from PLA-based adhesion barriers elevate FDG uptake, falsely mimicking cancer recurrence^63,64^.

Surprisingly, 2DG did not significantly reduce macrophage recruitment as measured by expression of CD11b or F4/80 in the PLA microenvironment. However, 2DG reduced macrophage activation into a proinflammatory phenotype (CD86), likely by competing with radiolabeled glucose for binding to hexokinase^36^, thereby inhibiting the first step in glycolysis. Since hydroxyapatite (HA) is often used to neutralize acidity from PLA degradation^10^, we compared effects of incorporating similar amounts (w/w) of HA to 2DG toward clinical translation of techniques targeting metabolism. Compared to 2DG, we observed increased pro-regenerative macrophage expression (CD206) with HA, which is consistent with the bioactivity of ceramic biomaterials^44^, opening the possibility of combinatorial strategies for regenerative applications. Corroborating CD206 results, our in-vitro data showed that neither 2DG nor 3PO, as a glycolytic inhibitor for PLA-based application, increases IL-4 or IL-10.

Compared to inhibiting glycolysis using 2DG, neutralizing acidity using HA increased macrophage recruitment and proinflammatory polarization, suggesting that metabolism and not acidity, is at the center of adverse immune responses to bulk PLA implants and PLA degradation products. Contrary to some studies, an explanation for the inability of HA to reduce inflammation in our study could be the amount used. Whereas the w/w concentration of HA present in our fabricated composites was 2 % for direct comparison to 2DG, greater than 20 % HA concentrations are more often used^65,66^. However, it is noteworthy that 2% HA resulted in significantly increased CD206 expression, suggesting pharmacological efficacy, yet could not reduce CD86 expression. Furthermore, unlike in soft tissue regeneration, enhanced mechanical properties of implants having more concentration of or comprising of only HA is desirable for bone tissue engineering^65,66^.

Increased fibroblast activation, measured by a-SMA expression, in the PLA microenvironment was reduced by inhibiting glycolysis using 2DG and not neutralizing acidity using HA. Compared to HA, 2DG reduced both a-SMA and TGF-β expression, suggesting that underlying metabolism regulates fibroblast activation in the PLA microenvironment. In agreement, metabolic reprogramming is known to play a key role in profibrotic disorders, activating fibroblasts^19^.

Most, if not all, publications on PLA’s biomedical application include a statement indicating that PLA breakdown products are metabolized through the tricarboxylic acid cycle. However, not until this study has it been demonstrated that bioenergetic changes occur in response to PLA. This key observation will redirect the field of tissue engineering, by offering an opportunity to intervene in this response. It opens up the possibilities to computationally identify relevant small molecules that could be clinically deployed, embedded in PLA implants, to mitigate adverse responses after carefully tuning drug release profiles. Moreover, use of PLA composites with ceramics could be optimized by combining the benefits of metabolic reprogramming with bioactivity of ceramics for bone tissue engineering. Beyond its ability to inhibit uptake of glycolytic substrates, related glutamine metabolic pathways, affected by aminooxyacetic acid could be explored for driving pro-regenerative macrophage response^67^.

Taken together, our findings suggest a model where PLA degradation products, including monomers of L-lactic acid, mechanistically remodel metabolism in cells of the immune microenvironment. This mechanism is specific and leads to increased proinflammatory cytokine and marker expression which can be modulated while stimulating anti-inflammatory cytokines. Our approach will enhance the biocompatibility and safety of biomaterials, including PLA-based implants for soft- and hard-tissue regeneration, significantly advancing tissue engineering.

## Methods

### Polylactide (PLA) materials and extraction

Highly crystalline PLA 3100HP and amorphous PLA 4060D (both from NatureWorks LLC) were used after their physicochemical and thermal properties were authenticated (Supplementary Table 1). PLA was sterilized by exposure to ultraviolet radiation for 30 minutes^25^. Afterwards, breakdown products (extracts)^21^ of PLA were obtained by suspending 4 g of PLA pellets in 25 ml of complete medium. Complete medium comprised of DMEM medium, 10% heat-inactivated Fetal Bovine Serum and 100 U/mL penicillin-streptomycin (all from ThermoFisher Scientific). PLA was extracted for 12 days in an orbital shaker at 250 rpm and 37 °C, after which extracts were decanted and extract’s pH measured. Either 100 or 150 *μ*l of extract (specified in each figure legend) was used per well of a flat-bottom 96-well plate; each volume was made up to 200 *μ*l, as final volume, using freshly made complete medium.

### Bioenergetic assessment

Bioluminescence was measured using the IVIS Spectrum in vivo imaging system (PerkinElmer) after adding 150 μg/mL of D-luciferin (PerkinElmer). Living Image (Version 4.5.2, PerkinElmer) was used for acquiring bioluminescence on the IVIS Spectrum. Standard ATP/ADP kits (Sigma-Aldrich) containing D-luciferin, luciferase and cell lysis buffer were used to according to manufacturer’s instructions. Luminescence at integration time of 1000 ms was obtained using the SpectraMax M3 Spectrophotometer (Molecular Devices) using SoftMax Pro (Version 7.0.2, Molecular Devices).

### pH measurements

The pH of extracts was assessed using an Orion Star A111 Benchtop pH Meter (ThermoFisher Scientific) under room temperature conditions (20 °C).

### Microscopy

Z-stack microscopy was accomplished by using the DeltaVision deconvolution imaging system (GE Healthcare) and softWoRx software (Version 7.2.1, GE Healthcare) at excitation and emission wavelengths of 525 and 558 nm, respectively for FITC. Section thickness of 0.2 μm for 64 to 128 sections were obtained at 40x and 100x magnification while imaging. Chambered Coverglass (Nunc Lab-Tek II) was used to seed 20,000 BGL cells (see cells below), keeping similar ratios as in a 96-well plate for volume of PLA extracts to volume of fresh medium.

### Glucose measurement

Glucose levels in complete medium was evaluated by a hand-held GM-100 glucose meter (BioReactor Sciences) after validation (Supplementary Fig. 1b) according to manufacturer’s instruction.

### Cells

Mouse embryonic fibroblast cell line (NIH 3T3 cell line; ATCC) and murine primary bone-marrow derived macrophages (BMDMs) were used. In each experiment, either 5,000 fibroblasts or 50,000 BMDMs were initially seeded. BMDMs were sourced from male and female C57BL/6J mice (Jackson Laboratories) of 3-4 months^30,35^. NIH 3T3 cells were stably transfected with a Sleeping Beauty transposon plasmid (pLuBIG) having a bidirectional promoter driving an improved firefly luciferase gene (fLuc) and a fusion gene encoding a Blasticidin-resistance marker (BsdR) linked to eGFP (BGL)^26^; enabling us to simultaneously monitor morphological and bioenergetic changes in live cells^68,69^. All cells were cultured in a total of 200 *μ*l complete medium with volumes of extracts specified in figure legends.

### Materials

3-(3-pyridinyl)-1-(4-pyridinyl)-2-propen-1-one (MilliporeSigma), 2-deoxyglucose (MilliporeSigma) and aminooxyacetic acid (Sigma-Aldrich) were used for glycolytic inhibition and L-lactic acid (Sigma-Aldrich) was used at various concentrations to reproduce the effects of PLA degradation products. Each of these materials were made in complete medium before adding to wells of a 96-well plate.

### Cell viability

Cell viability was assessed using the crystal violet staining assay^29^, at room temperature, as an end-point measure of total biomass generated over the course of the culture period. Briefly, out of 200 *μ*l of medium per well, 150 *μ*l is discarded. To each well, 150 *μ*l of 99.9 % methanol (MilliporeSigma) is added for 15 mins to kill and fix the cells, then discarded. Afterwards, 100 *μ*l of 0.5 % crystal violet (25 % methanol) is added for 20 mins, then the wells are emptied. Each well is washed twice with 200 *μ*l of phosphate buffered saline for 2 mins. Absorbance (optical density) was acquired at 570 nm using the the SpectraMax M3 Spectrophotometer (Molecular Devices) and SoftMax Pro software (Version 7.0.2, Molecular Devices).

### Functional metabolism

Basal measurements of oxygen consumption rate (OCR), extracellular acidification rate (ECAR) and lactate-linked proton efflux rate (PER) were obtained in real-time using the Seahorse XFe-96 Extracellular Flux Analyzer (Agilent Technologies)^34–36^. Prior to running the assay, cell culture medium was washed with and replaced by the Seahorse XF DMEM medium (pH 7.4) supplemented with 25 mM D-glucose and 4 mM Glutamine. The Seahorse plates were equilibrated in a non-CO2 incubator for an hour prior to the assay. The Seahorse ATP rate and cell energy phenotype assays were run according to manufacturer’s instruction and all reagents for the Seahorse assays were sourced from Agilent Technologies. Wave software (Version 2.6.1) was used to export Seahorse data directly as means ± standard deviation (SD).

### Chemokine and cytokine measurements

Cytokine and chemokine levels were measured using a MILLIPLEX MAP mouse magnetic bead multiplex kit (MilliporeSigma)^39^ to assess for IL-6, MCP-1, TNF-α, IL-1β, IL-4, IL-10, IFN-l and 1L-13 protein expression in supernatants. Data was acquired using Luminex 200 (Luminex Corporation) by the xPONENT software (Version 3.1, Luminex Corporation). Using the glycolytic inhibitor, 3PO, expectedly decreased cytokine values to < 3.2 pg/ mL in some experiments. For statistical analyses, those values were expressed as 3.1 pg/ mL. Values exceeding the dynamic range of the assay, in accordance with manufacturer’s instruction, were excluded. Additionally, IL-6 ELISA kits (RayBiotech) for supernatants were used according to manufacturer’s instructions.

### D/L-lactic acid determination assays

Measurements of L- and D-lactic acid were using standard D- and L-lactate assay kits (Sigma-Aldrich) according to manufacturer’s instruction after optimization (Supplementary Fig. 5a-b). Negative absorbance values which were outside the dynamic range for the assay were excluded during analysis.

### Optical rotation

Polarimetry was used to characterize the L-content and optical purity of the PLA samples with a P-2000 polarimeter (Jasco) by the Spectra Manager software (Version 2.13.00, Jasco). The optical rotation, [*α*]_25_, was measured and averaged for three samples of each polymer in chloroform (Omnisolv), at a concentration of 1 g/ mL. Conditions were set at 25 °C and 589 nm wavelength. Sucrose was used as a standard reference material, and its specific optical rotation was reported as approximately 67 °.

### Gel permeation chromatography

Gel permeation chromatography (GPC) was conducted to characterize the polymer molecular weights using a 600 controller (Waters) equipped with Optilab T-rEX refractive index (RI) and TREOS II multi-angle light scattering (MALS) detectors (Wyatt Technology Corporation), and a PLgel 5*μ*m MIXED-C column (Agilent Technologies) with chloroform eluent (1 mL/min). ASTRA software (Version 7.3.2.21, Wyatt Technology Corporation) was used. Polystyrene standards (Alfa Aesar) with Mn ranging from 35,000 to 900,000 Da were used for calibration.

### Differential scanning calorimetry

Differential scanning calorimetry (DSC) was conducted with a DSC Q20 (TA Instruments) to analyse the melting temperature (T_m_), glass transition temperature (T_g_), and percent crystallinity of the PLA grades. Thermal Advantage software (Version 5.5.23, TA Instruments) was used. Temperature was first equilibrated to 0 °C, then ramped up to 200 °C at a heating rate of 10 °C/ min; temperature was then held isothermally for 5 minutes. Afterwards, the sample was cooled back to 0 °C at a rate of 10 °C/min, then held isothermally for 2 minutes. Finally, the material was heated back to 200 °C at 10 °C/ min.

### In-vivo studies, tissue processing and analyses

Amorphous PLA was compounded with 2DG at 190 °C for 3 mins in a DSM 15 cc mini-extruder (DSM Xplore) and pelletizer (Leistritz Extrusion Technology). Our in-vitro studies indicate 1 mM 2DG to be an effective concentration. Accordingly, we estimated that 189 mg of 2DG in 10 g of amorphous PLA will approximate effective concentrations after accounting for potential thermal degradation of 2DG, converting mM to w/ w values^70^. We compounded comparable amounts (200 mg) of hydroxyapatite (HA; 2.5 *μ*m^2^ particle sizes^42^; Sigma-Aldrich) in 10 g of amorphous PLA under the same melt-blending thermal conditions. To exclude the effect of melt-blending as a confounder in our studies, amorphous PLA controls were processed under the same thermal conditions to make “reprocessed” amorphous PLA. Pellets from melt-blending were made into 1.75 mm diameter filaments using an extruder (Filabot EX2) at 170 °C with air set at 93. For surgical implantation, amorphous PLA filaments were cut into 1 mm lengths; four biomaterials were subcutaneously implanted on the dorsum (back) of each mouse, with two cranially (2.5 cm apart) and two caudally (2.5 cm apart)^12^.

Two-month old female C57BL/6J mice (n = 3 mice per group) with an average weight of 19 g were used according to procedures approved by the Institutional Animal Care and Use Committee at Michigan State University (PROTO202100327). Mice were anesthetized using isoflurane (2-3 %). The back of each mouse was shaved and alternate iodine and alcohol swabs were used as skin disinfectants. Aseptic surgery consisted of incisions through the skin into the subcutis, where biomaterials were inserted into a pouch made with forceps. Afterwards, surgical glue (3M Vetbond) was used to appose the skin. Each mouse received intraperitoneal or subcutaneous pre- and post-operative meloxicam (5 mg/ kg) injections as well as postoperative saline. Sham controls underwent the same procedure without biomaterial implantation. After 6 weeks, the dorsum of mice was shaved to visibly observe sites of surgical implantation. Thereafter, mice were intraperitoneally injected with 4.82 MBq F-18 fluorodeoxyglucose (Cardinal Health) in 200 *μ*l. At 65 mins post-dose, mice were euthanized and blood drawn from their hearts. Circular biopsies (12 mm diameter) of full skin thickness, with visible implants in the center, were recovered. Similar sized biopsies were collected from mice in the sham group in the region where surgical incision was made. Biomaterial migration from subcutaneous sites only allowed for the recovery of most and not all implants. As such, for obtaining data on the gamma counter (Fig. 7a), there were 12 skin biopsies from 3 mice in the sham group, 8 skin biopsies from 3 mice (amorphous PLA group) and 10 skin biopsies from 3 mice (amorphous + 2DG group). Skin biopsies, blood sample and heart organs were weighed, with only skin samples fixed in 4% paraformaldehyde (PFA). Activity in all samples was assessed via gamma counter (Wizard 2, Perkin Elmer) once decayed to a linear range. All injected doses and gamma counter measurements were decay-corrected to the same timepoint to calculate the percent of injected dose taken up per gram of assessed tissue (% ID / g; Fig. 7a).

For tissue staining, one skin biopsy per mouse was passed through increasing concentration of 10 %, 20 % and 30 % sucrose, daily. Using 99.9% methanol (Sigma-Aldrich) on dry ice, tissues were embedded in optimal cutting temperature (O.C.T.) compound (Tissue-Tek) by snap freezing. After equilibration at −20 °C, multiple successive 8 *μ*m sections were obtained using a microtome-cryostat. Sections were routinely stained using hematoxylin and eosin. Two different tissue sections were immunostained using conjugated antibodies as follows: 1) F4/80-FITC (1:100; BioLegend; 123107), CD11b-PE (1:100; BioLegend; 101207), CD206-BV421 (1:200; BioLegend; 141717) and CD86-Alexa Fluor 647 (1:100; BioLegend; 105019) using ordinary mounting medium; 2) alpha-SMA-eFluor660 (1:150; ThermoFisher Scientific; 50-9760-82), TGF-beta-PE (1:100; ThermoFisher Scientific; 12-9821-82) using DAPI mounting medium. Sections for TGF-beta were permeabilized using 0.1% Triton X in 1x PBS (PBST) for 8 mins then washed off with 1x PBS generously. Afterwards, blocking buffer (0.5 % bovine serum albumin in 1x PBS) was used to cover slides for 30 mins. Slides were then incubated in antibodies at 4 °C overnight. Subsequently, slides with tissue sections were washed in 1x PBS, and mounting medium applied.

Immunostained sections on slides were imaged using a Leica DMi8 Thunder microscope fitted with a DFC9000 GTC sCMOS camera and LAS-X software (Leica, version 3.7.4). Imaging settings at 20x magnification and 100 % intensity were: 1) F4/80-FITC excitation using the 475 laser (filter 535/ 70; 500 ms); CD11b-PE excitation using the 555 laser (no filter; 500 ms); CD206-BV421 excitation using 395 laser (no filter; 150 ms); CD86-Alexa Fluor 647 excitation using the 635 laser (no filter; 500 ms). 2) alpha-SMA-eFluor660 excitation using the 635 laser (no filter; 500 ms), TGF-beta-PE excitation using the 555 laser (no filter; 500 ms) and DAPI excitation using the 395 laser (535 filter; 500 ms). On the other hand, sections stained with hematoxylin and eosin were imaged at 40x using the Nikon Eclipse Ci microscope fitted with a CoolSNAP DYNO (Photometrics) and NIS elements BR 5.21.02 software (Nikon Instruments Inc.). Microscope images were prepared and analyzed using ImageJ (version 1.53k). For analyzing immunostained sections, 5 randomly selected rectangular areas of interest (1644.708 *μ*m^2^), encompassing cells adjacent to implants, were obtained as mean gray values^71^ a tissue section. In the sham group, biopsies were taken from incision sites and areas without cells were also analyzed. Where derived from n = 2 or n = 3 mice, 10 or 15 data points, respectively are graphically represented to fully reveal inherent variance across samples (Fig. 7b-e; Fig. 8a-b); only the aPLA + HA group had sections derived from n = 2 mice after one sample was damaged during cryo-sectioning and excluded from analyses. Representative images (16-bit; 0 to 65,535) were adjusted to enhance contrast for direct comparison using ImageJ as follows: CD86 (800 – 11,000), CD206 (2,000 – 5,000), F4/80 (500 – 4,000), CD11b (800 – 11,000), SMA (1,300 – 5,000), DAPI (6,000 – 31, 000), TGF (1,900 – 13,000).

### Statistics and reproducibility

Statistical software (GraphPad Prism) was used to analyse data presented as mean with standard deviation (SD). Significance level was set at p < 0.05, and details of statistical tests and sample sizes, which are biological replicates, are provided in figure legends. Exported data (mean, SD) from Wave in Seahorse experiments had the underlying assumption of normality and similar variance, and thus were tested using corresponding parametric tests as indicated in figure legends.

## Data availability

The data supporting the findings of this study are available within the paper and its Supplementary Information.

## Acknowledgements

Euthanized C57BL/6J mice were a gift from RR Neubig (facilitated by J Leipprandt) and the Campus Animal Resources at Michigan State University (MSU). Funding for this work was provided in part by the James and Kathleen Cornelius Endowment at MSU.

## Author contributions

Conceptualization, C.V.M. and C.H.C.; Methodology, C.V.M., K.R.Z., K.D.H., S.B.G., R.N. and C.H.C.; Investigation, C.V.M., M.A., E.U., M.O.B., M.M.K., K.S., A.V.M., H.P., S.C., J.M.H, C.L.M., S.J.C., M.H. and A.T.; Writing – Original Draft, C.V.M.; Writing – Review & Editing, C.V.M., M.A., E.U., M.O.B., M.M.K., K.S., A.V.M., H.P., S.C., J.M.H, C.L.M., S.J.C., M.H., A.T., K.R.Z., K.D.H., S.B.G., R. N. and C.H.C.; Funding Acquisition, C.H.C.; Resources, R.N. and C.H.C.; Supervision, K.R.Z., K.D.H., S. B.G., R.N. and C.H.C.

## Competing interests

C.V.M and C.H.C are inventors on a pending patent application filed by Michigan State University on metabolic reprogramming to biodegradable polymers.

## References

1 Farah, S., Anderson, D. G. & Langer, R. Physical and mechanical properties of PLA, and their functions in widespread applications - A comprehensive review. Adv Drug Deliv Rev 107, 367–392, doi:10.1016/j.addr.2016.06.012 (2016).

2 Givissis, P. K., Stavridis, S. I., Papagelopoulos, P. J., Antonarakos, P. D. & Christodoulou, A. G. Delayed foreign-body reaction to absorbable implants in metacarpal fracture treatment. Clinical Orthopaedics and Related Research® 468, 3377–3383 (2010).

3 Laine, P., Kontio, R., Lindqvist, C. & Suuronen, R. Are there any complications with bioabsorbable fixation devices?: a 10 year review in orthognathic surgery. International journal of oral and maxillofacial surgery 33, 240–244 (2004).

4 Chalidis, B., Kitridis, D., Savvidis, P., Papalois, A. & Givissis, P. Does the Inion OTPStm absorbable plating system induce higher foreign-body reaction than titanium implants? An experimental randomized comparative study in rabbits. Biomedical Materials 15, 065011 (2020).

5 Poh, P. S. et al. Polylactides in additive biomanufacturing. Advanced Drug Delivery Reviews 107, 228–246 (2016).

6 Ramot, Y., Haim-Zada, M., Domb, A. J. & Nyska, A. Biocompatibility and safety of PLA and its copolymers. Adv Drug Deliv Rev 107, 153–162, doi:10.1016/j.addr.2016.03.012 (2016).

7 Mosier-LaClair, S., Pike, H. & Pomeroy, G. Intraosseous bioabsorbable poly-L-lactic acid screw presenting as a late foreign-body reaction: a case report. Foot & ankle international 22, 247–251 (2001).

8 Athanasiou, K. A., Agrawal, C. M., Barber, F. A. & Burkhart, S. S. Orthopaedic applications for PLA-PGA biodegradable polymers. Arthroscopy 14, 726–737, doi:10.1016/s0749-8063(98)70099-4 (1998).

9 Waris, E. et al. Long-term bone tissue reaction to polyethylene oxide/polybutylene terephthalate copolymer (Polyactive®) in metacarpophalangeal joint reconstruction. Biomaterials 29, 2509–2515 (2008).

10 Agrawal, C. M. & Athanasiou, K. A. Technique to control pH in vicinity of biodegrading PLA-PGA implants. J Biomed Mater Res 38, 105–114, doi:10.1002/(sici)1097-4636(199722)38:2<105::aid-jbm4>3.0.co;2-u (1997).

11 Taylor, M. S., Daniels, A. U., Andriano, K. P. & Heller, J. Six bioabsorbable polymers: in vitro acute toxicity of accumulated degradation products. J Appl Biomater 5, 151–157, doi:10.1002/jab.770050208 (1994).

12 Deng, M. et al. Dipeptide-based Polyphosphazene and Polyester Blends for Bone Tissue Engineering. Biomaterials 31, 4898–4908, doi:10.1016/j.biomaterials.2010.02.058 (2010).

13 Pajares-Chamorro, N. et al. Silver-doped bioactive glass particles for in vivo bone tissue regeneration and enhanced methicillin-resistant Staphylococcus aureus (MRSA) inhibition. Materials Science and Engineering: C 120, 111693 (2021).

14 Lih, E. et al. A Bioinspired Scaffold with Anti-Inflammatory Magnesium Hydroxide and Decellularized Extracellular Matrix for Renal Tissue Regeneration. ACS Cent Sci 5, 458–467, doi:10.1021/acscentsci.8b00812 (2019).

15 Xu, T. O., Kim, H. S., Stahl, T. & Nukavarapu, S. P. Self-neutralizing PLGA/magnesium composites as novel biomaterials for tissue engineering. Biomed Mater 13, 035013, doi:10.1088/1748-605X/aaaa29 (2018).

16 Kamata, M., Sakamoto, Y. & Kishi, K. Foreign-body reaction to bioabsorbable plate and screw in craniofacial surgery. Journal of Craniofacial Surgery 30, e34–e36 (2019).

17 Narayanan, G., Vernekar, V. N., Kuyinu, E. L. & Laurencin, C. T. Poly (lactic acid)-based biomaterials for orthopaedic regenerative engineering. Advanced drug delivery reviews 107, 247–276 (2016).

18 Gonzalez-Lomas, G., Cassilly, R. T., Remotti, F. & Levine, W. N. Is the etiology of pretibial cyst formation after absorbable interference screw use related to a foreign body reaction? Clinical Orthopaedics and Related Research® 469, 1082–1088 (2011).

19 Xie, N. et al. Glycolytic reprogramming in myofibroblast differentiation and lung fibrosis. American journal of respiratory and critical care medicine 192, 1462–1474 (2015).

20 O’Neill, L. A. & Pearce, E. J. in J Exp Med Vol. 213 15–23 (2016).

21 Ignatius, A. A. & Claes, L. E. In vitro biocompatibility of bioresorbable polymers: poly(L, DL-lactide) and poly(L-lactide-co-glycolide). Biomaterials 17, 831–839, doi:10.1016/0142-9612(96)81421-9 (1996).

22 Yang, Y. et al. In vitro degradation of porous poly (l-lactide-co-glycolide)/β-tricalcium phosphate (PLGA/β-TCP) scaffolds under dynamic and static conditions. Polymer Degradation and Stability 93, 1838–1845 (2008).

23 Bostman, O. M. & Pihlajamaki, H. K. Adverse tissue reactions to bioabsorbable fixation devices. Clin Orthop Relat Res, 216–227 (2000).

24 Choueka, J. et al. Canine bone response to tyrosine-derived polycarbonates and poly(L-lactic acid). J Biomed Mater Res 31, 35–41, doi:10.1002/(sici)1097-4636(199605)31:1<35::aid-jbm5>3.0.co;2-r (1996).

25 Athanasiou, K. A., Niederauer, G. G. & Agrawal, C. M. Sterilization, toxicity, biocompatibility and clinical applications of polylactic acid/polyglycolic acid copolymers. Biomaterials 17, 93–102, doi:10.1016/0142-9612(96)85754-1 (1996).

26 Kanada, M. et al. Differential fates of biomolecules delivered to target cells via extracellular vesicles. Proceedings of the National Academy of Sciences 112, E1433–E1442 (2015).

27 Frottin, F. et al. The nucleolus functions as a phase-separated protein quality control compartment. Science 365, 342–347 (2019).

28 Hui, S. et al. Glucose feeds the TCA cycle via circulating lactate. Nature 551, 115–118 (2017).

29 Feoktistova, M., Geserick, P. & Leverkus, M. Crystal violet assay for determining viability of cultured cells. Cold Spring Harbor Protocols 2016, pdb. prot087379 (2016).

30 Gonçalves, R. & Mosser, D. M. The isolation and characterization of murine macrophages. Current protocols in immunology 111, 14.11.11–14.11.16 (2015).

31 Infantino, V., Iacobazzi, V., Palmieri, F. & Menga, A. ATP-citrate lyase is essential for macrophage inflammatory response. Biochemical and biophysical research communications 440, 105–111 (2013).

32 Jijon, H. B. et al. Inhibition of poly (ADP-ribose) polymerase attenuates inflammation in a model of chronic colitis. American Journal of Physiology-Gastrointestinal and Liver Physiology 279, G641–G651 (2000).

33 Tan, Z. et al. in The Journal of biological chemistry Vol. 290 46–55 (2015).

34 Ip, W. E., Hoshi, N., Shouval, D. S., Snapper, S. & Medzhitov, R. Anti-inflammatory effect of IL-10 mediated by metabolic reprogramming of macrophages. Science 356, 513–519 (2017).

35 Mills, E. L. et al. Succinate Dehydrogenase Supports Metabolic Repurposing of Mitochondria to Drive Inflammatory Macrophages. Cell 167, 457–470.e413, doi:10.1016/j.cell.2016.08.064 (2016).

36 Tannahill, G. et al. Succinate is a danger signal that induces IL-1β via HIF-1α. Nature 496, 238–242, doi:10.1038/nature11986 (2013).

37 Clem, B. et al. Small-molecule inhibition of 6-phosphofructo-2-kinase activity suppresses glycolytic flux and tumor growth. Molecular cancer therapeutics 7, 110–120 (2008).

38 Kauppinen, R. A., Sihra, T. S. & Nicholls, D. G. Aminooxyacetic acid inhibits the malate-aspartate shuttle in isolated nerve terminals and prevents the mitochondria from utilizing glycolytic substrates. Biochim Biophys Acta 930, 173–178, doi:10.1016/0167-4889(87)90029-2 (1987).

39 Sprague, L. et al. Dendritic cells: in vitro culture in two-and three-dimensional collagen systems and expression of collagen receptors in tumors and atherosclerotic microenvironments. Experimental cell research 323, 7–27 (2014).

40 Philpott, G. W. et al. RadioimmunoPET: detection of colorectal carcinoma with positron-emitting copper-64-labeled monoclonal antibody. Journal of Nuclear Medicine 36, 1818–1824 (1995).

41 Wen, S.-S. et al. Metabolic reprogramming and its clinical application in thyroid cancer. Oncology Letters 18, 1579–1584 (2019).

42 Pérez, E. Mechanical performance of in vitro degraded polylactic acid/hydroxyapatite composites. Journal of Materials Science 56, 19915–19935 (2021).

43 Graney, P. L., Lurier, E. B. & Spiller, K. L. Biomaterials and bioactive factor delivery systems for the control of macrophage activation in regenerative medicine. ACS Biomaterials Science & Engineering 4, 1137–1148 (2017).

44 Xue, H. et al. Enhanced tissue regeneration through immunomodulation of angiogenesis and osteogenesis with a multifaceted nanohybrid modified bioactive scaffold. Bioactive Materials (2022).

45 Veiseh, O. et al. Size- and shape-dependent foreign body immune response to materials implanted in rodents and non-human primates. Nat Mater 14, 643–651, doi:10.1038/nmat4290 (2015).

46 Pariente, J.-L., Kim, B.-S. & Atala, A. In vitro biocompatibility evaluation of naturally derived and synthetic biomaterials using normal human bladder smooth muscle cells. The Journal of urology 167, 1867–1871 (2002).

47 Andersson, S. R., Hakkarainen, M., Inkinen, S., Södergård, A. & Albertsson, A.-C. Polylactide stereocomplexation leads to higher hydrolytic stability but more acidic hydrolysis product pattern. Biomacromolecules 11, 1067–1073 (2010).

48 Kozlov, A. M., Lone, A., Betts, D. H. & Cumming, R. C. Lactate preconditioning promotes a HIF-1α-mediated metabolic shift from OXPHOS to glycolysis in normal human diploid fibroblasts. Scientific reports 10, 1–16 (2020).

49 Olive, A. J., Kiritsy, M. & Sassetti, C. (Am Assoc Immnol, 2021).

50 Iliopoulos, D., Hirsch, H. A. & Struhl, K. An epigenetic switch involving NF-κB, Lin28, Let-7 MicroRNA, and IL6 links inflammation to cell transformation. Cell 139, 693–706 (2009).

51 Eming, S. A., Wynn, T. A. & Martin, P. Inflammation and metabolism in tissue repair and regeneration. Science 356, 1026–1030 (2017).

52 Jiang, W. & Xu, J. Immune modulation by mesenchymal stem cells. Cell proliferation 53, e12712 (2020).

53 Pajarinen, J. et al. Mesenchymal stem cell-macrophage crosstalk and bone healing. Biomaterials 196, 80–89 (2019).

54 Swartzlander, M. D. et al. Immunomodulation by mesenchymal stem cells combats the foreign body response to cell-laden synthetic hydrogels. Biomaterials 41, 79–88 (2015).

55 Van den Bossche, J. et al. Mitochondrial dysfunction prevents repolarization of inflammatory macrophages. Cell reports 17, 684–696 (2016).

56 Manoharan, I., Prasad, P. D., Thangaraju, M. & Manicassamy, S. Lactate-Dependent Regulation of Immune Responses by Dendritic Cells and Macrophages. Frontiers in Immunology, 3062 (2021).

57 Zhang, A. et al. Lactate-induced M2 polarization of tumor-associated macrophages promotes the invasion of pituitary adenoma by secreting CCL17. Theranostics 11, 3839 (2021).

58 Lin, S. et al. Lactate-activated macrophages induced aerobic glycolysis and epithelial-mesenchymal transition in breast cancer by regulation of CCL5-CCR5 axis: a positive metabolic feedback loop. Oncotarget 8, 110426 (2017).

59 Romero-Garcia, S., Moreno-Altamirano, M. M. B., Prado-Garcia, H. & Sánchez-García, F. J. Lactate contribution to the tumor microenvironment: mechanisms, effects on immune cells and therapeutic relevance. Frontiers in immunology 7, 52 (2016).

60 Samuvel, D. J., Sundararaj, K. P., Nareika, A., Lopes-Virella, M. F. & Huang, Y. Lactate boosts TLR4 signaling and NF-κB pathway-mediated gene transcription in macrophages via monocarboxylate transporters and MD-2 up-regulation. The Journal of Immunology 182, 2476–2484 (2009).

61 Yang, K. et al. Lactate Suppresses Macrophage Pro-Inflammatory Response to LPS Stimulation by Inhibition of YAP and NF-κB Activation via GPR81-Mediated Signaling. Frontiers in Immunology 11, 2610 (2020).

62 Tang, C.-Y. & Mauro, C. Similarities in the metabolic reprogramming of immune system and endothelium. Frontiers in immunology 8, 837 (2017).

63 Hsieh, T.-C. & Hsu, C.-W. Foreign body reaction mimicking local recurrence from polyactide adhesion barrier film after laparoscopic colorectal cancer surgery: A retrospective cohort study. Medicine 101(2022).

64 Chong, G. O., Lee, Y. H., Hong, D. G., Cho, Y. L. & Lee, Y. S. Unabsorbed polylactide adhesion barrier mimicking recurrence of gynecologic malignant diseases with increased 18F-FDG uptake on PET/CT. Archives of Gynecology and Obstetrics 292, 191–195 (2015).

65 Bernardo, M. P. et al. PLA/Hydroxyapatite scaffolds exhibit in vitro immunological inertness and promote robust osteogenic differentiation of human mesenchymal stem cells without osteogenic stimuli. Scientific reports 12, 1–15 (2022).

66 Kim, S.-S., Park, M. S., Jeon, O., Choi, C. Y. & Kim, B.-S. Poly (lactide-co-glycolide)/hydroxyapatite composite scaffolds for bone tissue engineering. Biomaterials 27, 1399–1409 (2006).

67 Jha, A. K. et al. Network integration of parallel metabolic and transcriptional data reveals metabolic modules that regulate macrophage polarization. Immunity 42, 419–430, doi:10.1016/j.immuni.2015.02.005 (2015).

68 Kanada, M. et al. Microvesicle-mediated delivery of minicircle DNA results in effective gene-directed enzyme prodrug cancer therapy. Molecular cancer therapeutics 18, 2331–2342 (2019).

69 Negrin, R. S. & Contag, C. H. In vivo imaging using bioluminescence: a tool for probing graft-versus-host disease. Nature Reviews Immunology 6, 484–490 (2006).

70 Poon, C. Measuring the density and viscosity of culture media for optimized computational fluid dynamics analysis of in vitro devices. Journal of the Mechanical Behavior of Biomedical Materials 126, 105024 (2022).

71 Bravo-Hernandez, M. et al. Spinal subpial delivery of AAV9 enables widespread gene silencing and blocks motoneuron degeneration in ALS. Nature medicine 26, 118–130 (2020).

