## Supplementary for "Polylactide Degradation Activates Immune Cells by Metabolic Reprogramming"

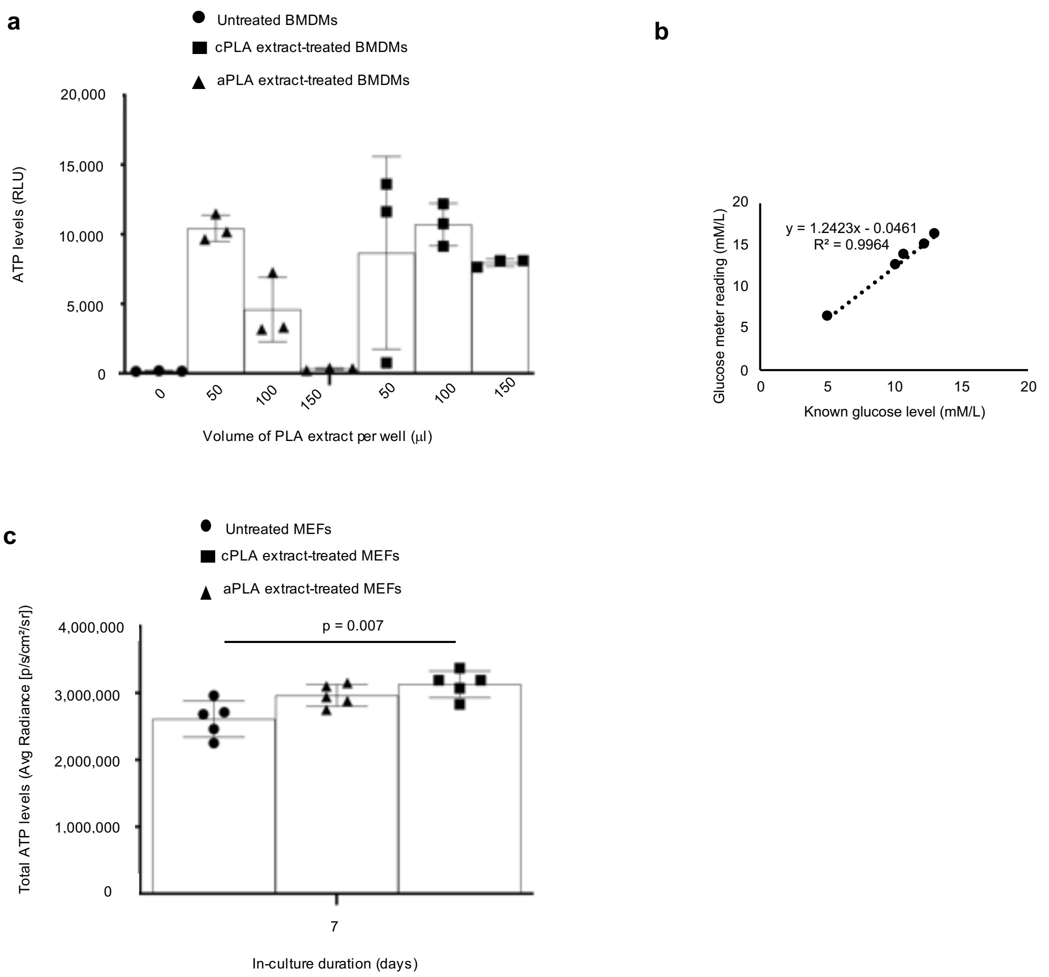


**Supplementary Figure 1 | Different doses of polylactide (PLA) extract alter bioenergetic (ATP) levels in primary bone marrow-derived macrophages (BMDMs) and using the glucose meter can measure glucose levels in cell culture medium.** **a**, Dose-bioenergetic response of the different PLA extracts on BMDMs revealed tendencies to alter ATP levels for all tested doses. **b**, Known glucose levels in cell culture medium linearly correlated (R square = 0.9964) with measurements from the glucose meter. **c**, Bioenergetic (ATP) levels are higher in mouse embryonic fibroblasts (MEFs) exposed to PLA extracts in comparison to controls. Mean (SD), n = 3 (Supplementary Fig. 1a), n = 5 (Supplementary Fig. 1c), simple linear regression, one-way ANOVA followed by Tukey's post-hoc test; crystalline PLA (cPLA), amorphous PLA (aPLA); 100 µl of control or PLA extract was used in Supplementary Fig. 1c.

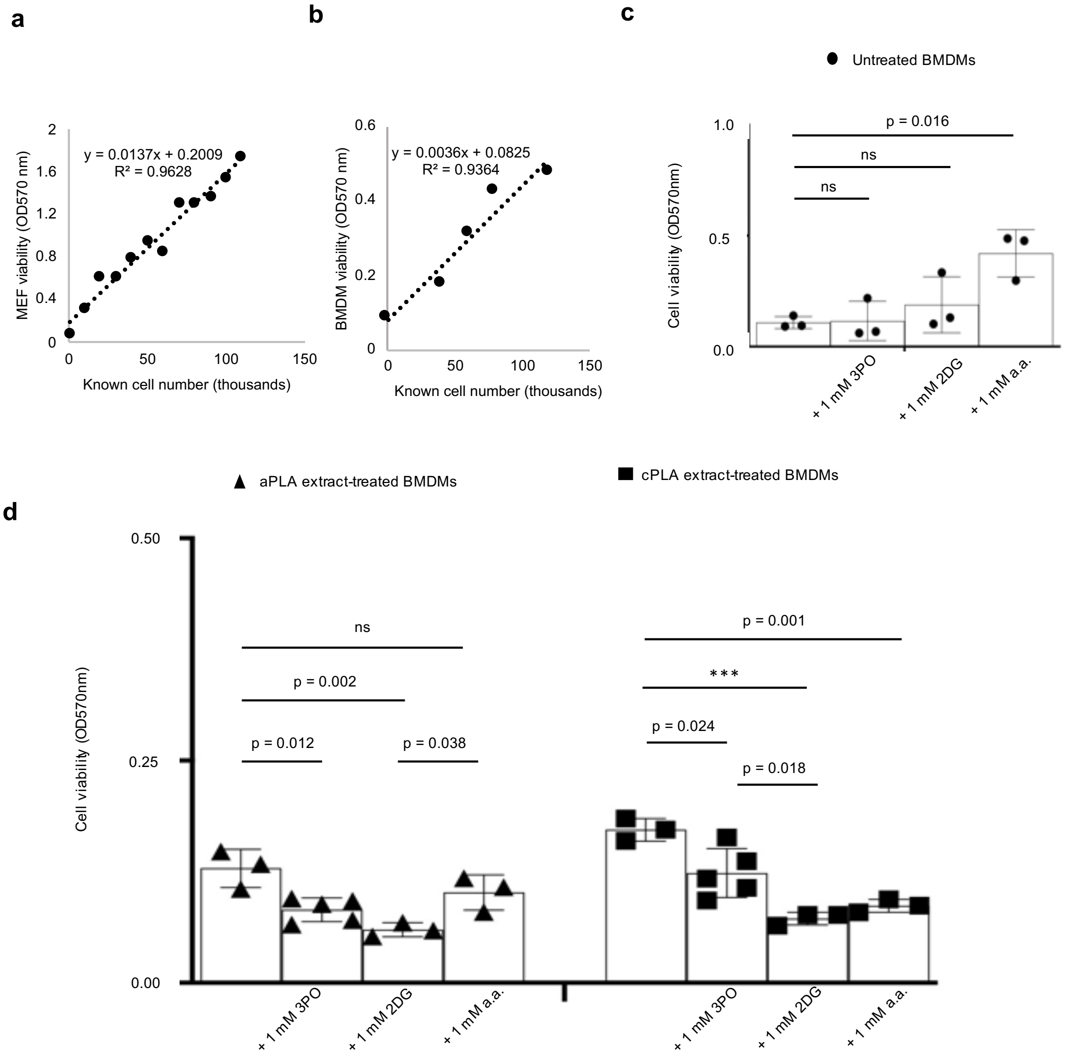


**Supplementary Figure 2 | Crystal violet assay can measure cell viability and cytotoxicity was selective for cells exposed to polylactide (PLA) following treatment with glycolytic inhibitors. a-b**, Known cell numbers linearly correlated with absorbance for **a**, mouse embryonic fibroblasts (MEF; R square = 0.9628) and **b**, primary bone marrow-derived macrophages (BMDMs; R square = 0.9364). **c-d**, Although cell viability was not decreased in untreated BMDMs following exposure to glycolytic inhibitors (**c**), BMDMs exposed to amorphous PLA (aPLA) or crystalline PLA (cPLA) degradation products (extract) decreased in cell viability after treatment with glycolytic inhibitors (**d**). Not significant (ns), *** $p < 0.001$, mean (SD), n = 3 (Supplementary Fig. 2c), n = 3-5 (Supplementary Fig. 2d), one-way ANOVA followed by Tukey's post-hoc test; 3-(3-pyridinyl)-1-(4-pyridinyl)-2-propen-1-one (3PO), 2-deoxyglucose (2DG) and aminooxyacetic acid (a.a.); 100 µl of control or PLA extract was used on day 7.

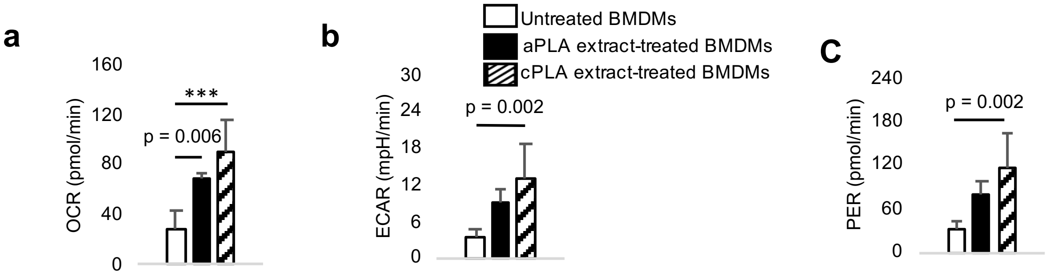


**Supplementary Figure 3 | Functional metabolic indices are increased in primary bone marrow-derived macrophages (BMDMs) after exposure to crystalline PLA (cPLA) degradation products (extracts). a-c**, Oxygen consumption rate (OCR, **a**), extracellular acidification rate (ECAR, **b**) and proton efflux rate (PER, **c**) are increased following exposure to cPLA extracts. *** $p < 0.001$, mean (SD), n=5, one-way ANOVA followed by Tukey's post-hoc test; 150 µl of control or PLA extract was used on day 7.

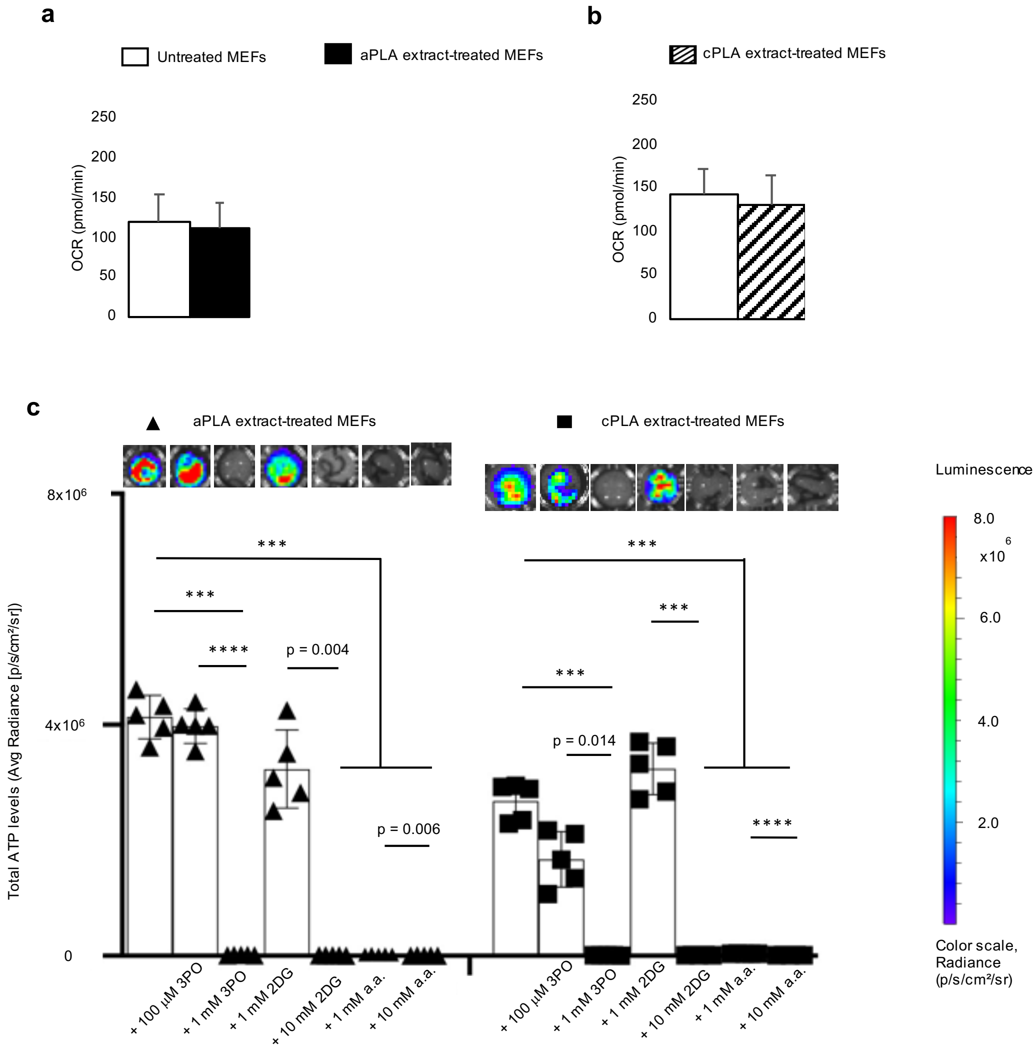


**Supplementary Figure 4 | Oxygen consumption rate (OCR) is not altered in mouse embryonic fibroblasts (MEFs) following prolonged exposure to polylactide (PLA) degradation products (extract)**. **a-b**, Following exposure to amorphous PLA (aPLA; **a**) or crystalline PLA (cPLA; **b**) extracts, OCR is unaffected. **c**, Bioenergetic levels on day 12 in MEFs exposed to aPLA or cPLA extracts are decreased in a dose-dependent manner by 3-(3-pyridinyl)-1-(4-pyridinyl)-2-propen-1-one (3PO), 2-deoxyglucose (2DG) and aminooxyacetic acid (a.a.). *** $p < 0.001$, **** $p < 0.0001$, mean (SD), n = 3 (Supplementary Fig. 4a, b), n = 5 (Supplementary Fig. 4c), two-tailed unpaired t-test or Brown-Forsythe and Welch ANOVA followed by Dunnett's T3 multiple comparisons test; 100 µl of control or PLA extract was used on day 7 (Supplementary Fig. 4a, b) or 12 (Supplementary Fig. 4c).

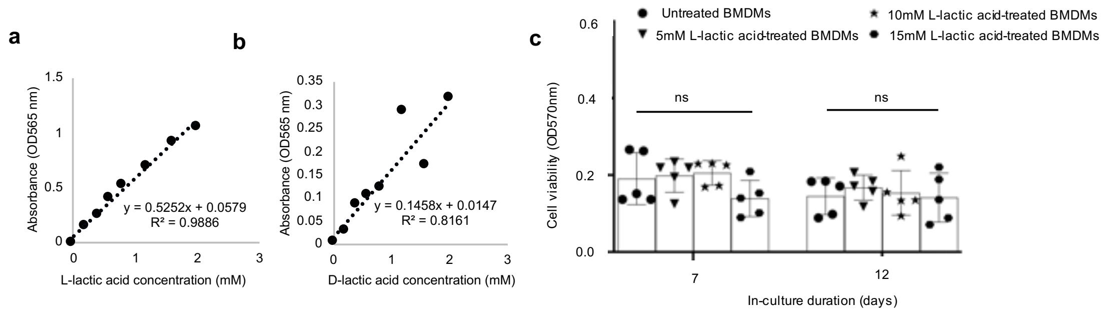


**Supplementary Figure 5 | D- and L-lactic acid levels can be detected by absorbance and cell viability is similar among macrophages treated with L-lactic acid. a-b**, Known concentrations of L-lactic (R square = 0.9886; **a**) and D-lactic (R square = 0.8161; **b**) acid linearly correlate with absorbance. **c**, Viability of primary bone marrow-derived macrophages (BMDMs) is similar after treatment with L-lactic acid over time. Not significant (ns), one-way ANOVA, mean (SD), n=5

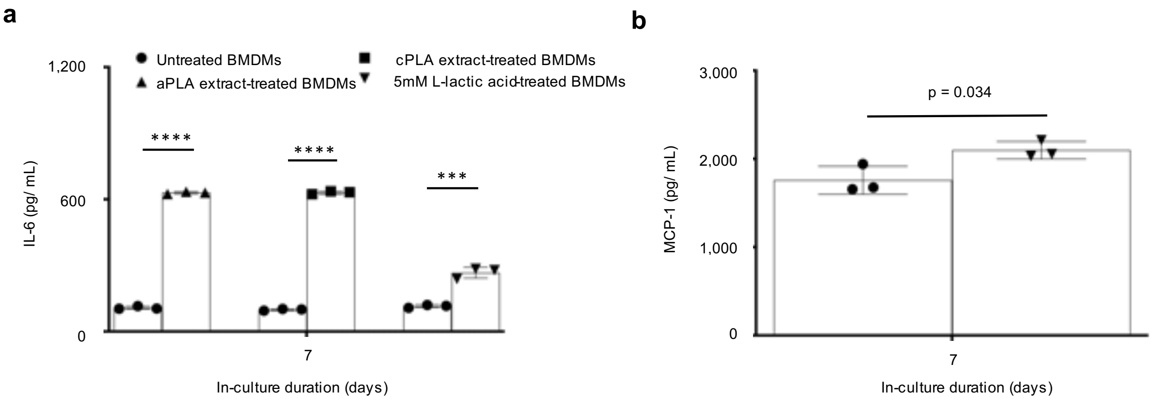


**Supplementary Figure 6 | IL-6 and MCP-1 protein levels are increased following prolonged exposure of primary bone marrow-derived macrophages (BMDMs) to L-lactic acid in comparison to untreated BMDMs. a,** Using ELISA reproduced changes in IL-6 levels following exposure of BMDMs to amorphous PLA (aPLA), crystalline PLA (cPLA) or L-lactic acid. **b**, Similarly, MCP-1 levels are increased after exposing BMDMs to L-lactic acid as measured by the MILLIPLEX assay. ***p<0.001, ****p<0.0001, mean (SD), n=3, two-tailed unpaired t-test; 100 µl of aPLA or 150 µl of cPLA with corresponding controls were used; whereas corresponding controls for PLA were incubated for 12 days, the controls for L-lactic acid were not.

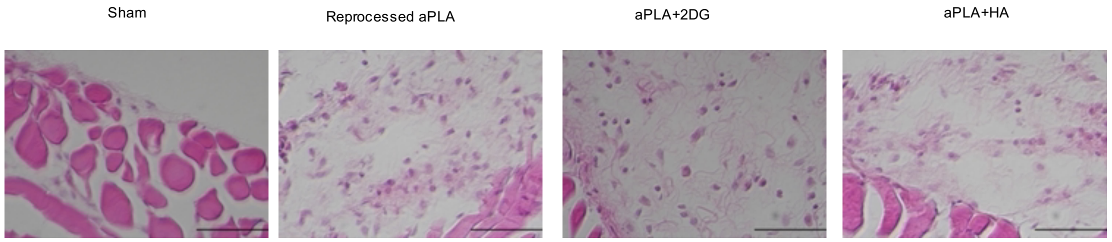


**Supplementary Figure 7 | Recruitment of inflammatory cellular infiltrates following implantation of amorphous polylactide (aPLA) with and without 2-deoxyglucose (2DG) or hydroxyapatite (HA) is compared to sham controls in cryo-sections stained using hematoxylin/ eosin (scale bars, 20 µm).**

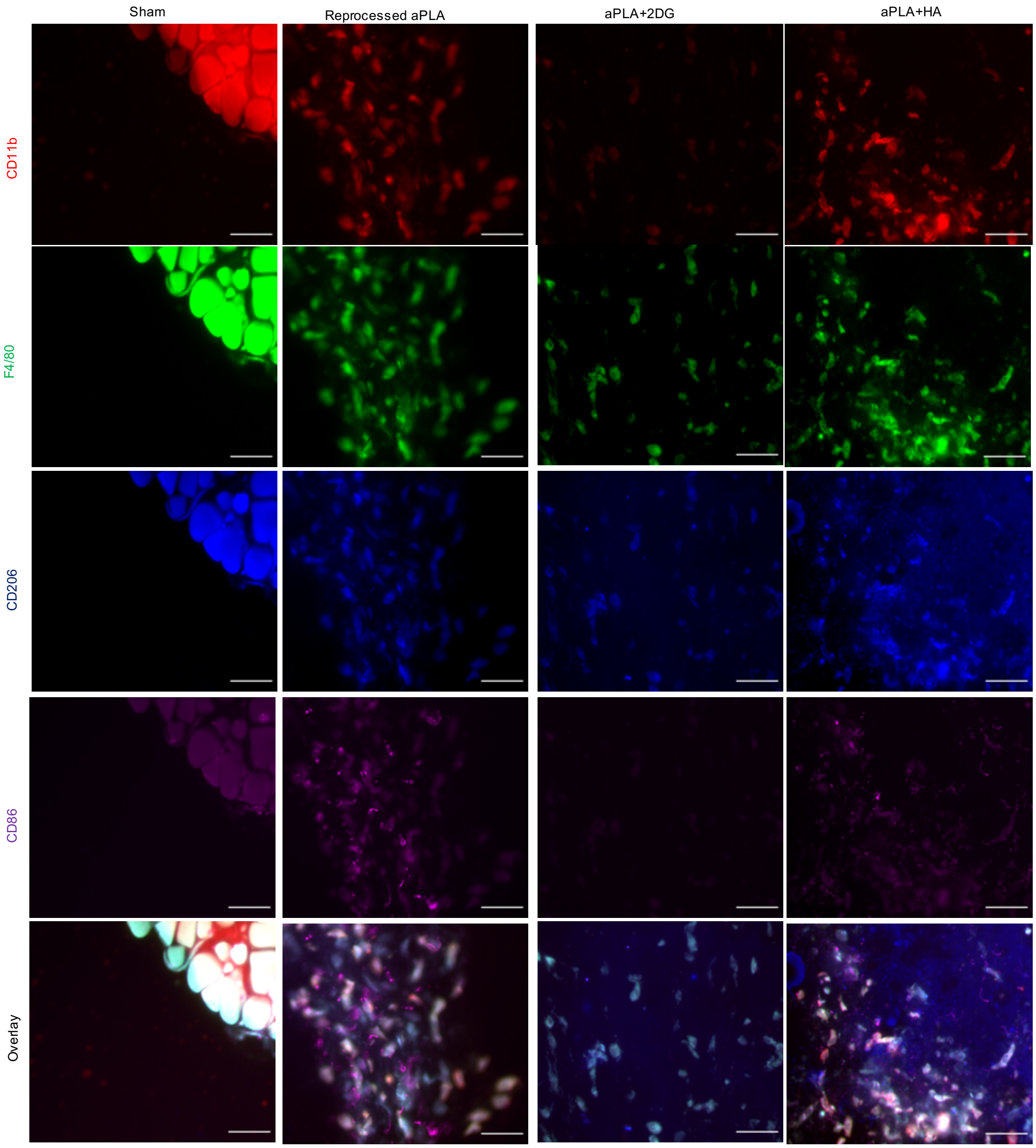


**Supplementary Figure 8 | Immunohistochemical staining with CD11b-PE, F4/80-FITC, CD206-BV421 and CD86-AF647 reveal the presence and polarization of macrophages following implantation of amorphous polylactide (aPLA) with and without 2-deoxyglucose (2DG) or hydroxyapatite (HA) when compared to sham controls (scale bars, 50 µm).**

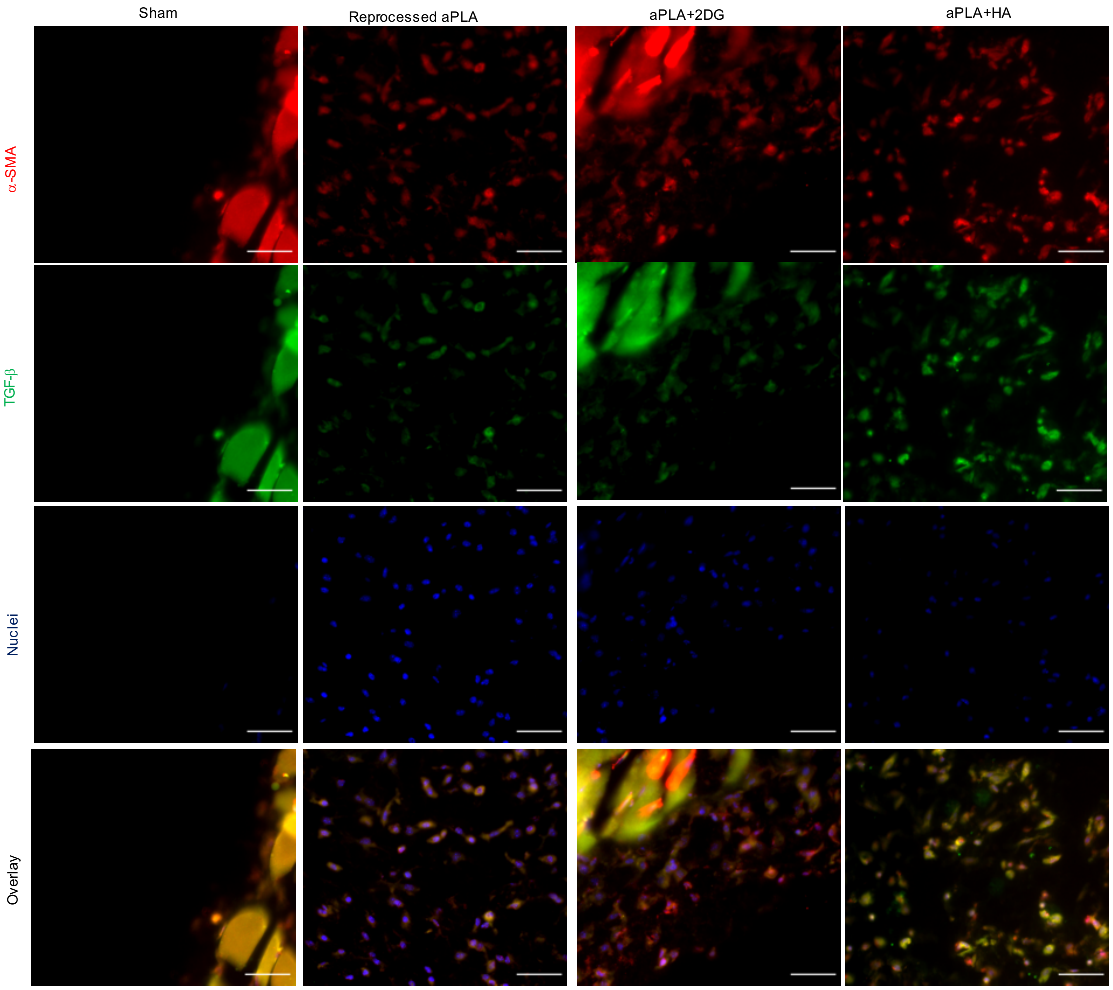


**Supplementary Figure 9 | Immunohistochemical staining with α-SMA-eFluor 660 and TGF-β-PE using a DAPI mounting medium show fibroblast activation following implantation of amorphous polylactide (aPLA) with and without 2-deoxyglucose (2DG) or hydroxyapatite (HA) when compared to sham controls (scale bars, 50 μm).**

**Supplementary Table 1: Authentication of physicochemical and thermal properties of commercial polylactide (PLA).**

| Criteria | PLA 3100HP (Crystalline PLA) | PLA 4060D (Amorphous PLA) |
|---|---|---|
| Optical purity (%) | 99.04 | 81.47 |
| L-content (%) | 99.40 | 90.54 |
| Glass transition temperature $T_g$ ($^oC$)‡ | 62.20 | 59.05 |
| Melting temperature $T_m$ ($^oC$)‡ | 175.85 | N/A |
| Crystallinity (pellet, %)† | 51.46 | 0 |
| Crystallinity (resin, %)‡ | 42.49 | 0 |
| Number average molecular weights $M_n$ (Da) | 87,390 | 113,270 |
| Weight average molecular weights $M_w$ (Da) | 157,060 | 200,200 |
| Polydispersity index | 1.797 | 1.767 |

†Percentage crystallinity of pellets was determined based on the first heating cycle.
‡Percentage crystallinity of the resin, $T_g$, and $T_m$ were determined based on the second heating cycle.
Molecular weights were based on a calibration curve of polystyrene standards.

**Supplementary Table 2: Molecular weights of polylactide (PLA) samples decrease after extraction in medium or water.**

| Extracted in: | | Initial molecular weight (Da) | | Final molecular weight (Da) | | Decrease (%) | |
|---|---|---|---|---|---|---|---|
| | PLA sample | $M_n$ | $M_w$ | $M_n$ | $M_w$ | $M_n$ | $M_w$ |
| Serum-containing medium | cPLA | 87,390 ± 2,840 | 157,060 ± 3,640 | 75,155 ± 1,340 | 140,540 ± 2,390 | 14.0 | 10.5 |
| | aPLA | 113,270 ± 1,880 | 200,200 ± 2,150 | 100,923 ± 3,380 | 185,365 ± 3,900 | 10.9 | 7.4 |
| Milli-Q water | cPLA | – | – | 75,155 ± 1,340 | 140,540 ± 2,390 | 14.0 | 10.5 |
| | aPLA | – | – | 103,302 ± 2,180 | 185,250 ± 3,560 | 8.8 | 7.5 |

Number average molecular weight ($M_n$) and weight average molecular weight ($M_w$) are expressed as mean (SD) and based on a calibration curve of polystyrene standards, n=3; crystalline PLA (cPLA), amorphous PLA (aPLA); Dashed line indicates that initial weights are the same for PLA, irrespective of whether PLA will be extracted in water or complete medium.

**Supplementary Table 3: Monomers of L- and D-lactic acid are detectable in extracts of polylactide.**

| | L-lactic acid (OD565nm) | D-lactic acid (OD565nm) |
|---|---|---|
| Crystalline PLA (cPLA) extract | 0.0034 ± 0.0025 | 0.0021 ± 0.0010 |
| Amorphous PLA (aPLA) extract | 0.0051 ± 0.0036 | 0.0023 ± 0.0002 |
| Control (milli-Q water) | 0.0007 ± 0.0004 | 0.0008 ± 0.0001 |

Lactic acid absorbance is expressed as mean (SD), n=2-3.
